# Integrating Earth Observation and Graph Theory to Evaluate Urban Green Spaces Connectivity Across European Capitals

**DOI:** 10.64898/2026.01.29.702234

**Authors:** Costanza Borghi, Saverio Francini, Leonardo Chiesi, Stefano Mancuso, Liubov Tupikina, Guido Caldarelli, Jacopo Moi, Elia Vangi, Giovanni D’Amico, Giuseppe De Luca, Gherardo Chirici

## Abstract

**Context:** As global urbanization intensifies, Urban Green Spaces (UGS) are pivotal for biodiversity conservation and climate change mitigation. However, comparative assessments of UGS spatial configuration and connectivity across diverse urban landscapes remain limited.

**Objectives:** This study aims to assess the spatial arrangement and connectivity of UGS across 28 European capital cities. Additionally, we evaluate how Network Science metrics derived from Graph Theory can complement traditional landscape ecology metrics to provide a more comprehensive understanding of UGS at a large scale.

**Methods:** We developed a European Urban Vegetation Map using Earth observation data to classify UGS at 10m resolution across the selected capitals. We then analyzed UGS connectivity for each city utilizing 40 traditional landscape metrics and a Graph-Theory-based approach.

**Results:** While traditional landscape metrics effectively quantified fragmentation, they often remain strongly correlated with total vegetation abundance. In contrast, Network Science metrics provided specific insights into UGS functional connectivity, distinguishing the quality of ecological links beyond spatial proximity. This integration allowed us to cluster European capitals into three distinct typologies: unconnected compact cities, large metropolises with complex peri-urban dynamics, and high-connectivity cities with robust networks. These findings demonstrate that graph-based indices effectively complement traditional metrics, highlighting that relying solely on green space percentage is insufficient for assessing the ecological resilience of urban environments.

**Conclusions:** These results underscore the relevance of Earth observation-based UGS assessment and demonstrate that graph-based landscape connectivity analysis outperforms simple abundance metrics. Therefore, effective assessment requires integrating structural metrics with graph-based connectivity to support resilient urban biodiversity.

## 1. Introduction

With the global population projected to reach about 10 billion in 2050 (Gu et al. 2021), almost 70% of the global population is expected to live in urban areas by the same year (United Nations 2018). Urban Green Spaces (UGS) provide a vast range of ecosystem services (Stroud et al. 2022): carbon and pollutants removal (Han et al. 2024; Du et al. 2025), temperature regulation (Duncan et al. 2019), biodiversity conservation (Afrifa et al. 2022), and benefits to human mental and physical health (Konijnendijk 2023). A standardized approach to assessing UGS is essential to support policy makers and urban planners to improve urban quality of life, and build biodiverse and resilient cities, urgently needed under climate change (Norton et al. 2015).

The European Union recognizes UGS in the Biodiversity Strategy for 2030 and within the objectives of the European Green Deal and the Sustainable Development Goals. However, their assessments often rely on an oversimplified indicator - the percentage of area covered by UGS (Maes et al. 2019). For example, the European Commission’s 2023 guidance for Nature Urban Plans lists UGS percentage as the primary indicator (followed by tree cover and protected area percentages), with no attention to spatial arrangement despite landscape ecology research showing its importance (Turner and Gardner 2015; Herath et al. 2024).

Monitoring the spatial arrangement of UGS in cities (Francini et al. 2024) is crucial for implementing nature-based solutions, limiting fragmentation and degradation (Huang et al. 2021), and assessing connectivity (Clauzel et al. 2018; Tarabon et al. 2022). Traditionally, this analysis implements landscape ecology metrics from categorical land-use/land-cover maps (Uuemaa et al. 2013; Das et al. 2022). Recently, Network Science based on mathematical Graph Theory has emerged as a powerful approach for assessing UGS arrangement and functional connectivity (Urban and Keitt 2001; Kong et al. 2010; Hashemi et al. 2024), modeling landscapes as networks of habitat patches (i.e., nodes) linked by potential movement paths (Galpern et al. 2011). Compared to traditional heterogeneity metrics, which are mainly focused on patch geometry rather than mosaic function (Uuemaa et al. 2009; Kupfer 2012), graph-based metrics offer a more functional connectivity analysis (de la Barra et al. 2022).

While some efforts have been made for small areas or limited ecosystems (Villegas et al. 2021, 2024), to our knowledge, large-scale studies comparing graph theory with traditional metrics for UGS are lacking, likely due to three main constraints.

First, high-resolution standardized vegetation maps for a relevant number of urban areas are scarce (Yan et al. 2018; Brandt et al. 2023). In this context, remote sensing has witnessed significant advancements in vegetation mapping, primarily due to improvements in temporal, spectral, and spatial resolution of available products (Dash and Ogutu 2016; Wulder et al. 2019; Fraucqueur et al. 2019). In Europe, the Copernicus program (Jutz and Milagro-Pérez 2020) provides Earth observation data from its constellation of several passive and active Earth Observation satellites, the “Sentinels”. In this context, the Copernicus Land Monitoring Service (CLMS) produces several thematic products and in situ data. Some of them are available wall-to-wall in Europe, such as the High-Resolution map of trees or grassland cover, while others are specifically developed for urban areas, such as the Urban Atlas products (Congedo et al. 2016). Second, processing spatial data over large areas is computationally demanding, though cloud platforms like Google Earth Engine mitigate this (Gorelick et al. 2017).

Finally, standardized methods to convert vegetation maps into graphs are limited. Recently, Graphab (Foltête et al. 2012, 2021) has emerged as an operational tool for computing global and local network metrics and has been applied to habitat suitability, planning evaluation, and fragmentation/connectivity studies (Kohler et al. 2016; Duflot et al. 2018; Kim et al. 2019; Kim and Kang 2022; Lumia et al. 2023).

In this paper, we aimed to assess the combined potential use of graph theory in combination with more traditional landscape ecology metrics for quantifying the spatial arrangements of UGS across European capital cities. In this scenario, the present study aims to quantitatively analyze whether the percentage area covered by UGS is informative enough for city ranking and urban planning, establishing it as a reference value for systematic comparison with other indicators.

To do so, we first created an innovative 10 m resolution European Urban Vegetation Map (EUVM) covering 28 European capital cities (EU-28), distinguishing three main vegetation categories: (i) trees, (ii) shrubs, and (iii) grassland. To address computational issues and improve its reproducibility, the process was implemented using the Google Earth Engine (Gorelick et al., 2017) cloud computing platform.

After successfully validating the EUVM against field surveys collected through the Land Use and Coverage Area Frame Survey (LUCAS) program, we used the Graphab-based R package “graph4lg” (Savary et al. 2021) to calculate ten graph theory-based metrics also used in Network Science. These metrics were then compared with 30 more traditional landscape metrics - such as landscape aggregation, patch area and edges, core areas, diversity, and patch shape - calculated using the software FRAGSTATS and the related R package “landscapemetrics” (Hesselbarth et al. 2019, 2024). Finally, the results were compared and discussed to assess the relevance of the different metrics in evaluating the spatial arrangement of UGS within the EU-28 capital cities that, through a cluster analysis, were classified according to their (i) city size, (ii) peri-urban forest presence, and (iii) spatial organization of UGS.

## 2. Materials and methods

### 2.1. Study area

The study was conducted for the EU-28, including the European Union capital cities and London (Figure 1). The definition of city boundaries is a critical issue since it directly impacts the extension of UGS and the related landscape ecology metrics. Here, we used the city boundaries adopted for the 2017 Eurostat Urban Audit data collection, the so-called “Eurobarometer”, as they were specifically developed to maximize the comparability of statistics across the very different spatial arrangements of European cities, towns, and Functional Urban Areas (European Commission, 2016). The city boundaries’ geographical source - hereinafter referred to as Urban Audit (URAU) - is available at https://ec.europa.eu/eurostat/web/gisco/geodata/statistical-units/urban-audit (last accessed on 05/22/2025).

**Fig. 1.**
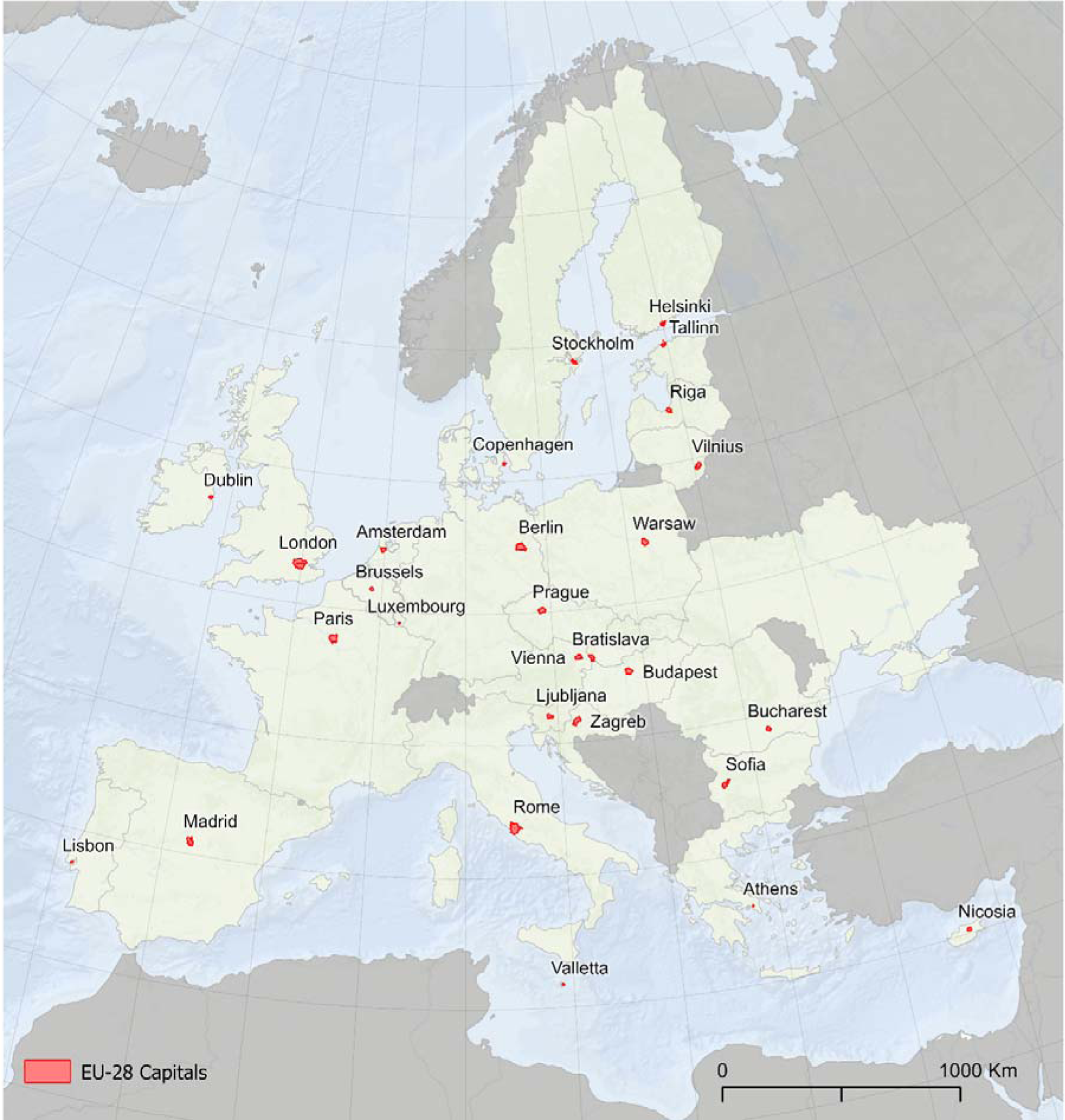
European capital cities investigated in the study, in red

Next, a 1 km buffer was created around each city to include vegetation patches that could fall across the URAU boundaries, but that are relevant for the ecological network connectivity analysis. Overall, we investigated an area of 11476.55 km^2^, covering approximately 0.3% of the total EU-28 area (447,687,200 ha). The cities investigated are very different in size, ranging from 1575 km^2^ in London to 39 km^2^ in Athens. Overall, the area hosts 52.4 million inhabitants, approximately 10% of the total population in the EU-28 region (Eurostat 2020).

### 2.2 The European Urban Vegetation Map

For the investigated area, we created a European Urban Vegetation Map (EUVM), with 10 m spatial resolution for the reference year 2018 and a nomenclature system based on three types of dominant vegetation cover: trees, shrubs, and herbaceous.

To create the EUVM, we innovatively aggregated four different Copernicus layers, including the high-resolution layers (HRL) with a spatial resolution of 5-10 m: (i) Urban Atlas (UA), (ii) Street Tree Layer (STL), (iii) HRL – Grassland (HRL-GRA), (iv) HRL – Tree Cover Density (HRL–TCD), and (v) HRL – Small Woody Features (HRL-SWF).

In brief, the Urban Atlas (UA) 2018 is a vector land use/land cover map having 17 urban classes with the Minimum Mapping Unit (MMU) of 0.25 ha and 10 rural classes with the MMU of 1 ha (ESA 2018a). Similarly, the Street Tree Layer (STL) – in our study reclassified as “trees”-is a vector layer aimed at mapping contiguous rows or patches of trees along the city streets that are too small for the MMU adopted in the UA; it has a MMU of 500 m^2^. Next, the HRL layers (i.e., GRA, TCD, SWF) were provided in raster format with a spatial resolution of 10 m. Currently, the 2018 reference year is available. In detail, HRL-GRA is a binary dataset distinguishing between grassland and non-grassland regions, as the result of a comprehensive approach that merges optical (Sentinel-2) and radar (Sentinel-1) data. For the 2018 reference year, GRA reported an overall accuracy is 95.31% (European Commission 2020). In this study, HRL-GRA was reclassified as “herbaceous”. Besides, HRL-TCD reports the level of tree cover density in a 0-100% range (ESA 2018b), based on Sentinel-2 data. For the 2018 reference year, the reported final overall accuracy of 96.43%. Here, the TCD HRL was reclassified into a Boolean map, considering those pixels with coverage equal to or above 10% as “trees” (1). The threshold was inherited from the internationally agreed-upon definition of forest area (FAO 2000). Similarly, the HRL-SWF is a raster Boolean layer with 5 m resolution mapping linear or patchy structures like hedgerows, trees, and shrubs along field or road margins, riparian vegetation, and scattered groups of trees and shrubs (EEA 2018), with an overall accuracy of 80% for the 2018 reference year. As for HRL-TCD, HRL-SWF was then reclassified as “trees”.

All the input layers were rasterized at 10 m resolution and projected in ETRS89 Lambert Azimuthal Equal Area (EPSG: 3035). To generate the EUVM, the data contained in the Copernicus layers were reclassified into four primary categories representing (0) non-vegetation, (1) trees, (2) shrubs, and (3) herbaceous vegetation, as in Table 1. The reclassification of the input maps and the further elaboration of the dataset at the European level were performed in the GEE environment (Gorelick et al. 2017), which allowed easier and faster computation.

**Table 1.**
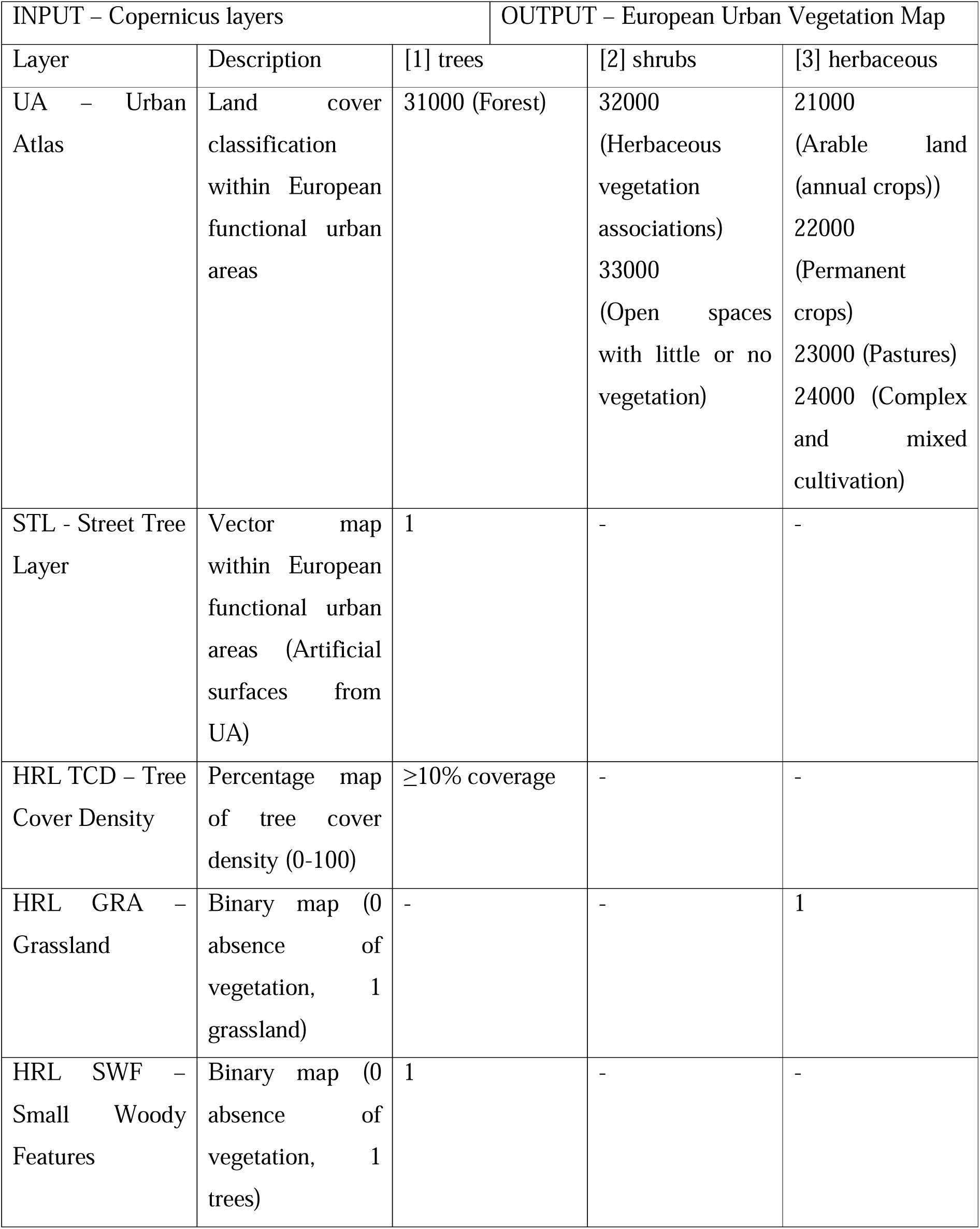
Reclassification scheme for Copernicus input layers. The codes not included are considered in the EUVM class (0), not vegetation.

The information in the four layers we considered is frequently redundant. For instance, most of the area classified as HRL-SWF has a tree cover from the HRL-TCD above the threshold of 10%, but not always. For this reason, in aggregating the layers for the same class, we adopted a conservative approach using the logical operator “OR” (Figure 2).

**Fig. 2.**
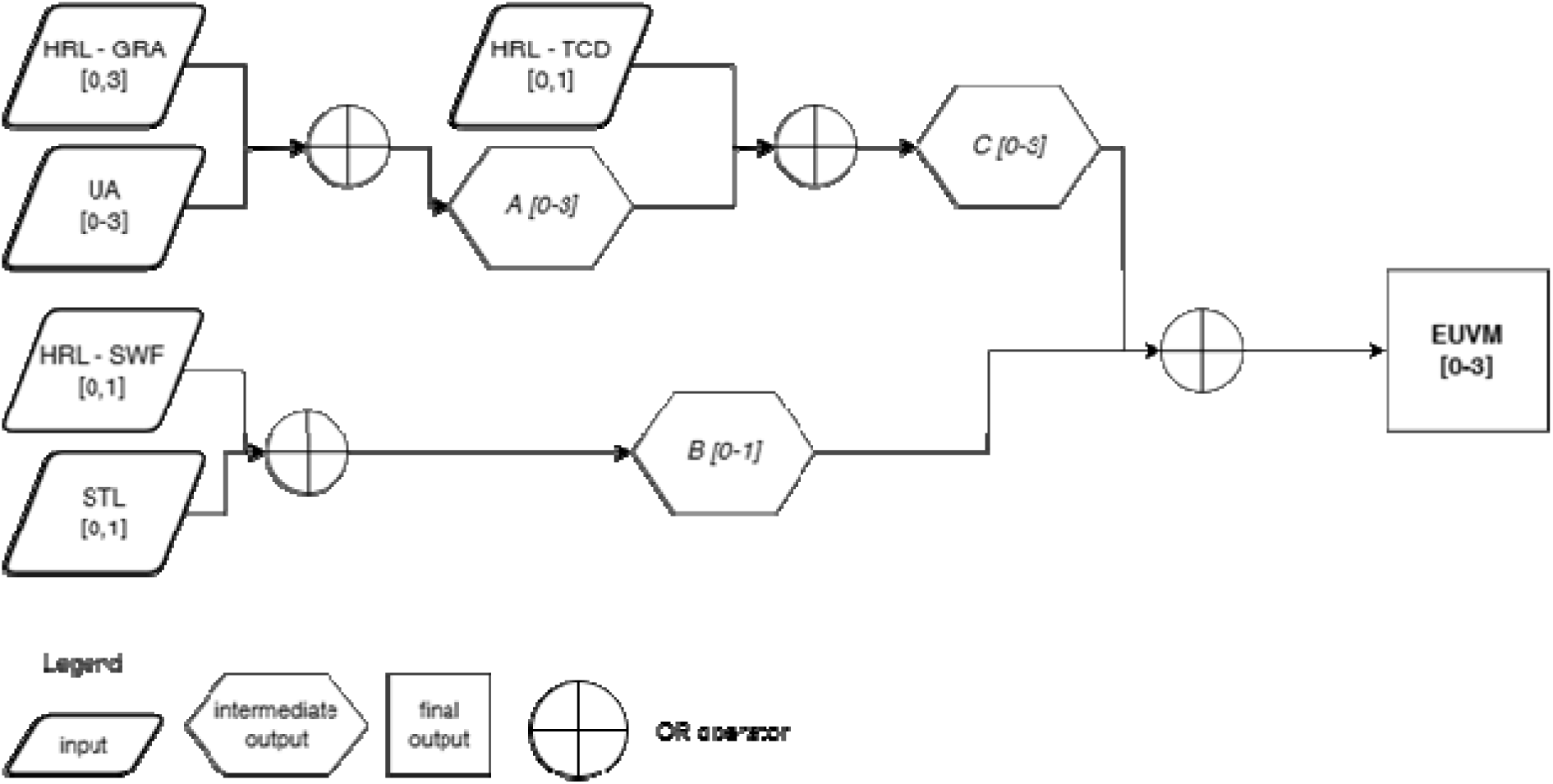
European Urban Vegetation Map flowchart

### 2.3 Data validation

Before adopting the EUVM for the following analyses, we validated it against the information reported in the Land Use and Coverage Area Frame Survey (LUCAS), for the reference year 2018. Starting in 2006, LUCAS has been conducted every three years to monitor land uses for social, economic, and ecological purposes based on a two-phase survey. In the first phase, a systematic grid of 2 km x 2 km across the EU (Gallego and Bamps 2008), (for a total of 1.1 million points) is photo interpreted with a system of nomenclature of 10 classes, in the second phase a subsample of the first phase sample is visited in the field (Ballin et al. 2018). The LUCAS database is openly available in GEE as a harmonized version of all yearly surveys (d’Andrimont et al. 2020).

For validating the EUVM, we used 337,845 LUCAS second-phase points visited in the field in 2018. Of these, 1866 points fall within the EU-28 city buffers where the EUVM was created. Due to the different definitions used in EUVM and LUCAS, we used a Boolean approach, considering together woodland/shrubland classes from both LUCAS and EUVM.

Based on the 2 x 2 confusion matrix from the LUCAS validation points, we obtained a true positive (TP) of 408, true negative (TN) of 1154, false positive (FP) 84, and false negative (FN) 220. Based on this, the resulting overall accuracy (OA) of the EUVM was 0.84, producer’s accuracy (PA) 0.64, and the user’s accuracy (UA) 0.80. Detailed city-level information is available in the Annex, Table 1A. On the basis of these results, we considered EUVM (Figure 3) as the basis for the following analysis.

**Fig. 3.**
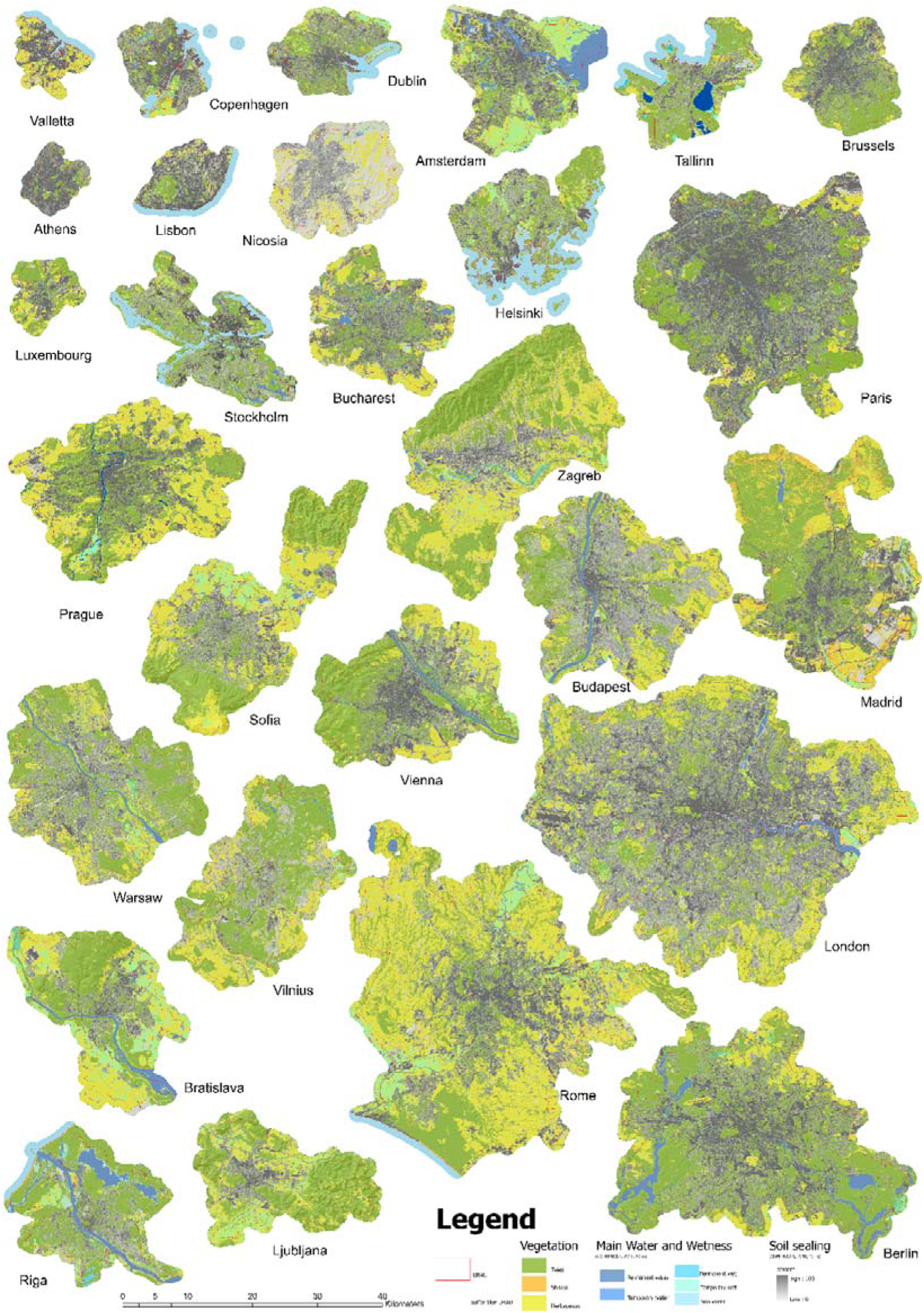
The EUVM of the 28 European capitals considered in this study. They are ranked from top left to bottom right based on the cluster analysis available in Figure 10. For a better comprehension of the maps we added to the layouts, the layers of the Copernicus High Resolution Layer for Water and Wetness and for Imperviousness

### 2.4 Network connectivity and landscape ecology analysis

For this study, two sets of landscape patterns and network connectivity metrics were computed using two different approaches. FRAGSTAT software (McGarigal et al. 2023) and the related R package “landscapemetrics” (Hesselbarth et al. 2024) were implemented for the calculation of the 30 landscape metrics related to the groups (i) Aggregation (AGG), (ii) Area and Edge (AED), (iii) Core Area (COR), (iv) Diversity (DIV) and (v) Shape (SHP) metrics. Besides, the software Graphab (Foltête et al. 2012, 2021) and the related R package “grap4lg” (Savary et al. 2021) were used for the calculation of the ten Network (NET) metrics. In both cases, the EUVM was used as the input of the analysis, setting the pixels classified as “trees” and “shrubs” as the target habitat categories of the network. The total list of 40 metrics we calculated for each city is reported in the Annex, Table 2A. The approach for the comparison of the indices was inspired by recent review (Uuemaa et al. 2009, 2013).

#### 2.4.1 Network connectivity metrics based on graph theory

For this approach, we used the Graphab metrics (Foltête et al. 2012, 2021) implemented in the R package “graph4lg” (Savary et al. 2021). First, the EUVM was used to create habitat patches when the cumulative area of adjacent pixels reached at least the minimum patch area threshold of 0.1 hectares (Bulman et al. 2007). Next, from each patch, a node is located in its centroid, and planar links are set between the habitat patches based on the Euclidean distance among them. Here, we consider the matrix as uniform in terms of resistance values of landscape categories. Lastly, the graph was pruned by setting a maximum distance threshold between patches of 200 m, which previous studies identify as suitable for urban forest landscape connectivity analysis (Chang-Fu et al. 2010).

Based on the graph, we calculated three global metrics that were considered relevant in recent studies on urban landscape connectivity analysis (e.g., Babí Almenar et al. 2019; Mu et al. 2021): (i) the probability of connectivity (PC), (ii) the (relative) equivalent connected area (ECA), and (iii) the integral index of connectivity (IIC).

The PC represents the probability that two individuals, randomly located in the landscape, are situated within interconnected habitat patches (Saura and Pascual-Hortal 2007). It is calculated as the area-weighted probability of connection between all patch pairs and is particularly sensitive to habitat loss. Next, ECA translates the PC value into an equivalent area of a single, maximally connected patch, facilitating interpretation in terms of habitat extent (Saura et al. 2011). To enable comparison across different cities, ECA values are standardized by the total area of each city (Savary et al. 2024). The IIC, in contrast, is based on a binary model of connectivity, reflecting the probability that individuals within a patch can reach each other through existing links. It provides a topological view of the network and increases with greater connectivity (Pascual-Hortal and Saura 2006). While PC relies on probabilistic interactions to capture species-specific movement across the landscape, IIC evaluates structural connectivity based on the presence or absence of links (Bodin and Saura 2010). Both indices range from 0 to 1, with higher values indicating better connectivity. In addition to these indices, eight additional metrics were used to describe the structure of the habitat network. These include the total, median, and Gini coefficient of patch capacity (area in hectares), the number of links per hectare (including those ≤200 m), their mean and standard deviation in distance, and the Gini index of link distances.

#### 2.4.2 Landscape ecology metrics

For this approach, we used FRAGSTAT metrics (McGarigal and Marks 1995; McGarigal et al. 2023) implemented in the R package “landscapemetrics” (Hesselbarth et al. 2024). Among the vast array of possible metrics, we selected the most relevant based on their use in previous experiences in urban landscapes (Kupfer 2012; Schindler et al. 2015; Chen and Yu 2017; Frazier and Kedron 2017; Das et al. 2020). Overall, we calculated a total of 30 metrics aggregated in five groups: i) aggregation (8), ii) area and edge (9), iii) core area (5), iv) diversity (2), and v) shape metrics (6). See Annex, Table 2A for a short description of the metrics and McGarigal et al. (2023) for a more complete explanation.

### 2.5. Comparing graph theory and traditional landscape ecology metrics

In this study, we employed a series of statistical analyses to explore the relationships among our metrics and their utility for urban classification and planning. First, we compared the correlation between the metrics, one against the other, reporting the Spearman’s coefficients for each pair. To understand the potential usefulness of the metrics for classifying the different cities according to specific metrics, we repeated the correlation analysis in two ways: first, based on rank scores assigned based on all the metrics (the largest values given the largest ranks), then using averaged metrics based on the six groups reported in Annex, Table 2A, which facilitate the interpretation of their relationship Next, to enhance the comprehension of our results and the potential for creating groups of cities with comparable conditions based on the calculated metrics, we performed two multivariate analysis: a hierarchical cluster analysis to formally identify clusters of cities based on the calculated metrics, and a Principal Component Analysis (PCA) of the original dataset, to highlight the relevance of the metrics in differentiating the spatial arrangement of UGS across cities.

Finally, one of the main goals of this study is to determine if the percentage area covered by UGS (specifically, *AED_pland*) is informative enough for city ranking and urban planning. Therefore, we used *AED_pland* as a reference value, systematically comparing it with all other indicators through correlation analysis across the cities.

## 3. Results

In more than half of the urban areas investigated (15 cities out of 28, Figure 4), the UGS share is higher than the European average (52.62%), based on the EUVM. Among those, Ljubljana has the largest share of UGS, covering 78.75% of its territory, mostly with trees (51.80%). On the opposite side, Athens showed the lowest UGS coverage, 19.46% of its territory, followed by Nicosia (25.96%) and Lisbon (26.53%).

**Fig. 4.**
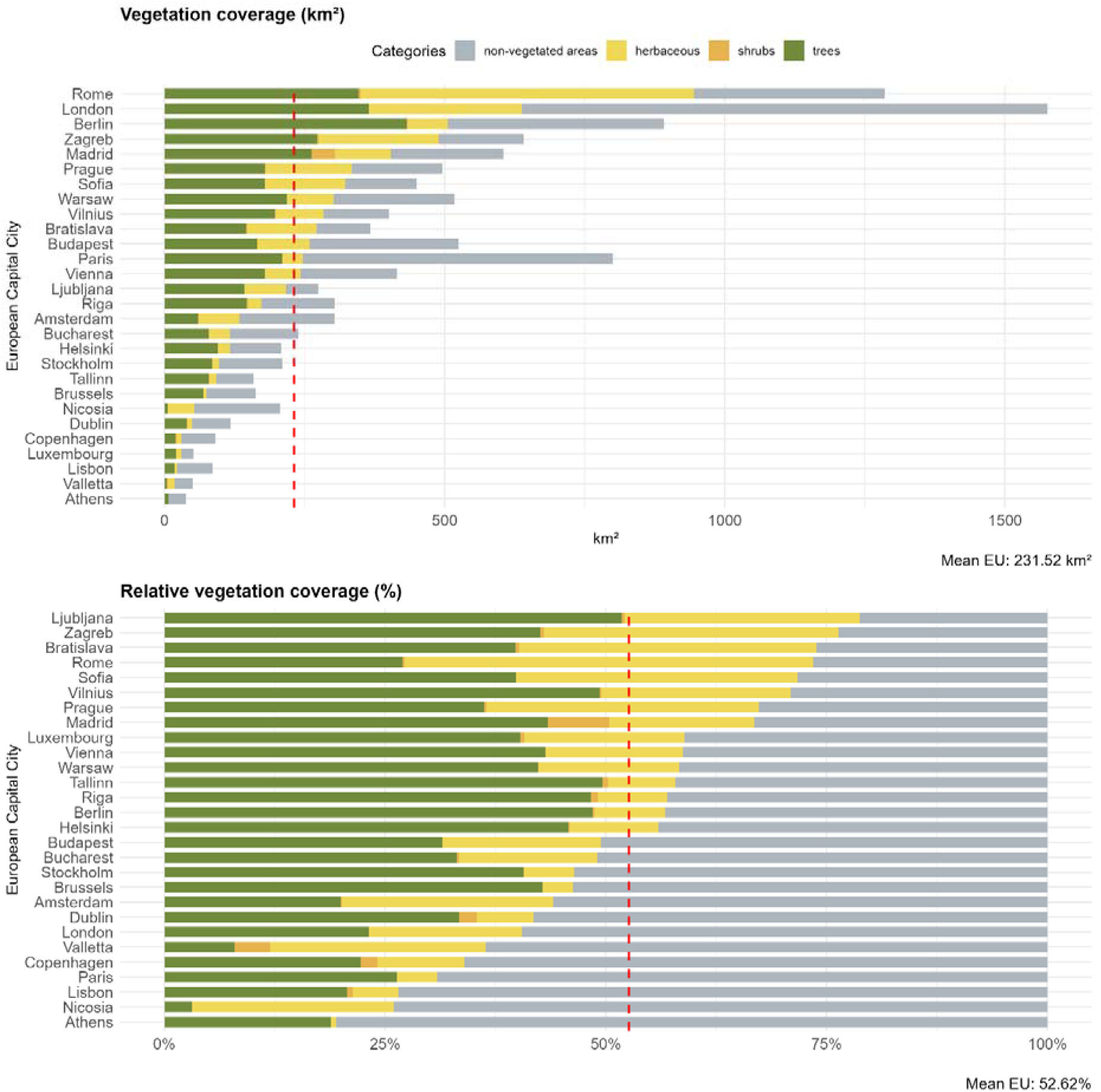
European capital cities’ land cover share in absolute values (above) and as a relative coverage (below) over the total area, according to the European Urban Vegetation Map. Cities are ranked based on the overall share of vegetation over the total area

Following Ljubljana, trees represent the most important UGS category in northern European areas, especially Tallinn and Vilnius (see Annex, Table 3A). In contrast, southern European capitals exhibit a significant proportion of herbaceous cover, notably in Rome, Bratislava, and Zagreb, along with shrubland, predominantly in Madrid and Valletta. Lastly, Mediterranean cities such as Athens recorded the lowest share of UGS among European capitals, with less than 20% vegetation coverage (19.45%). This is followed by Nicosia and Lisbon, each with approximately 26% vegetation coverage.

Based on these results, it is possible to note a clear relationship between the population density (i.e., number of inhabitants per area) and the per-capita availability of UGS. So, the higher the population density, the lower (with an exponential law) is the amount of UGS available per capita (Figure 5).

**Fig. 5.**
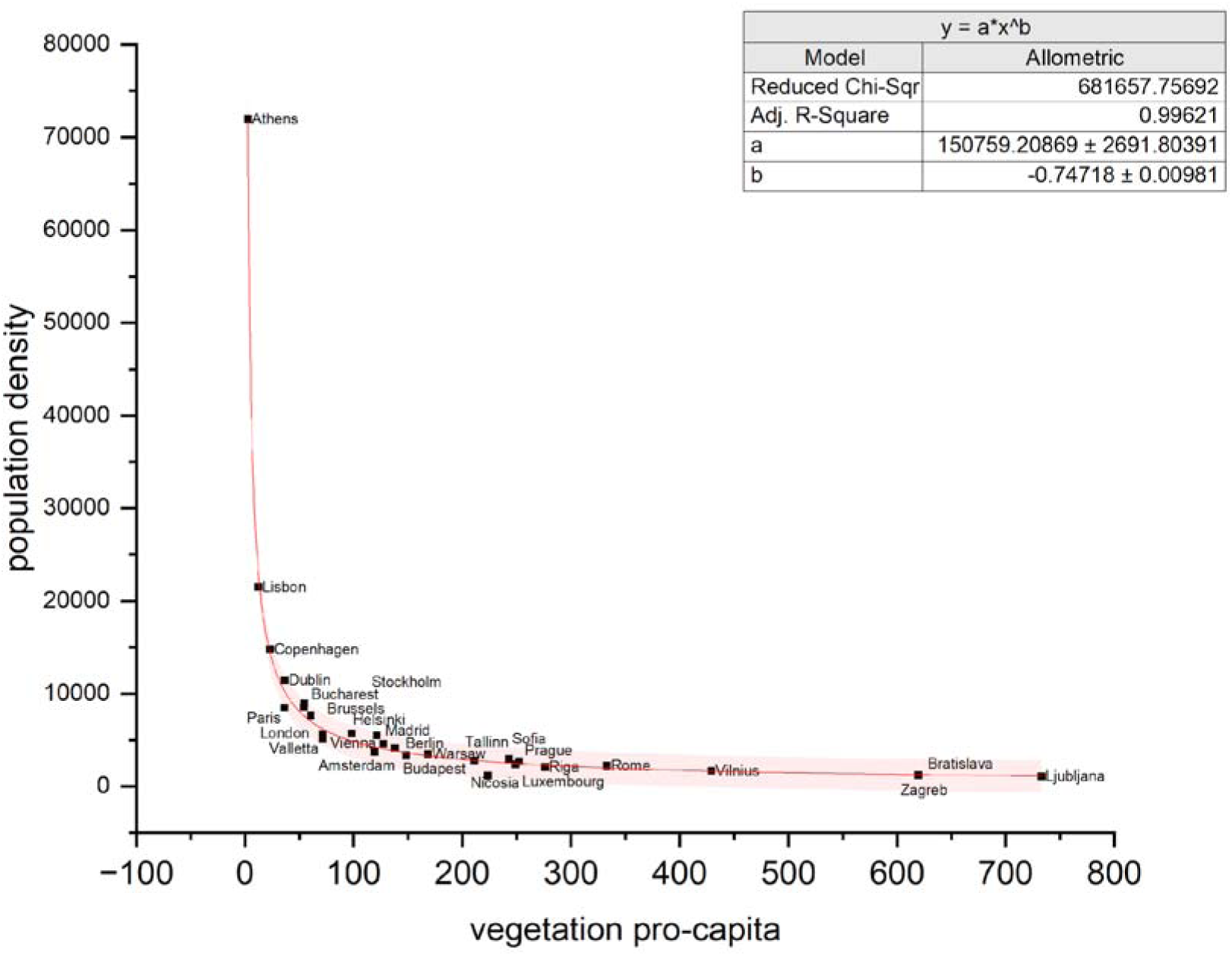
The relationship between the population density (inhabitants per square kilometers) versus the vegetation per capita (UGS area per inhabitant in square meters)

### 3.1 Landscape metrics

For each city, the average value of each landscape metric calculated is reported in the Annex, Tables 4-8A, while below we report the details for each group.

#### 3.1.1 Aggregation metrics

Several spatial aggregation metrics (AGG) are related to the percentage amount of UGS available in the different cities, but in general, the average relationship is quite weak (average R^2^ = 0.3). We found direct relationships with *AI, cohesion*, *enn_CV* or inverse relationships with *division*, *split* (with R^2^ up to 0.7). In other cases, the aggregation metrics are not related to the relative amount of UGS in the area and, for this reason, are probably more informative. This is the case of *np* and *LSI* that account respectively for the number of patches and their level of aggregation: a very good example is London where the UGS covers only the 23% of the total area (the 6^th^ lower value within the 28 capitals) but the number of patches (more than 27000) is instead the highest and very disaggregated resulting in the highest value of *LSI*. In general, *np* and *LSI* resulted, as expected, strictly related to each other with a direct linear relationship (R^2^ = 0.85).

#### 3.1.2 Area and edge metrics

Within the metrics related to patch area and edge (AED), some theoretical relationships are respected, such as the decreasing exponential relationship between the patch mean area and the number of patches per hectare of UGS (Figure 6).

**Fig. 6.**
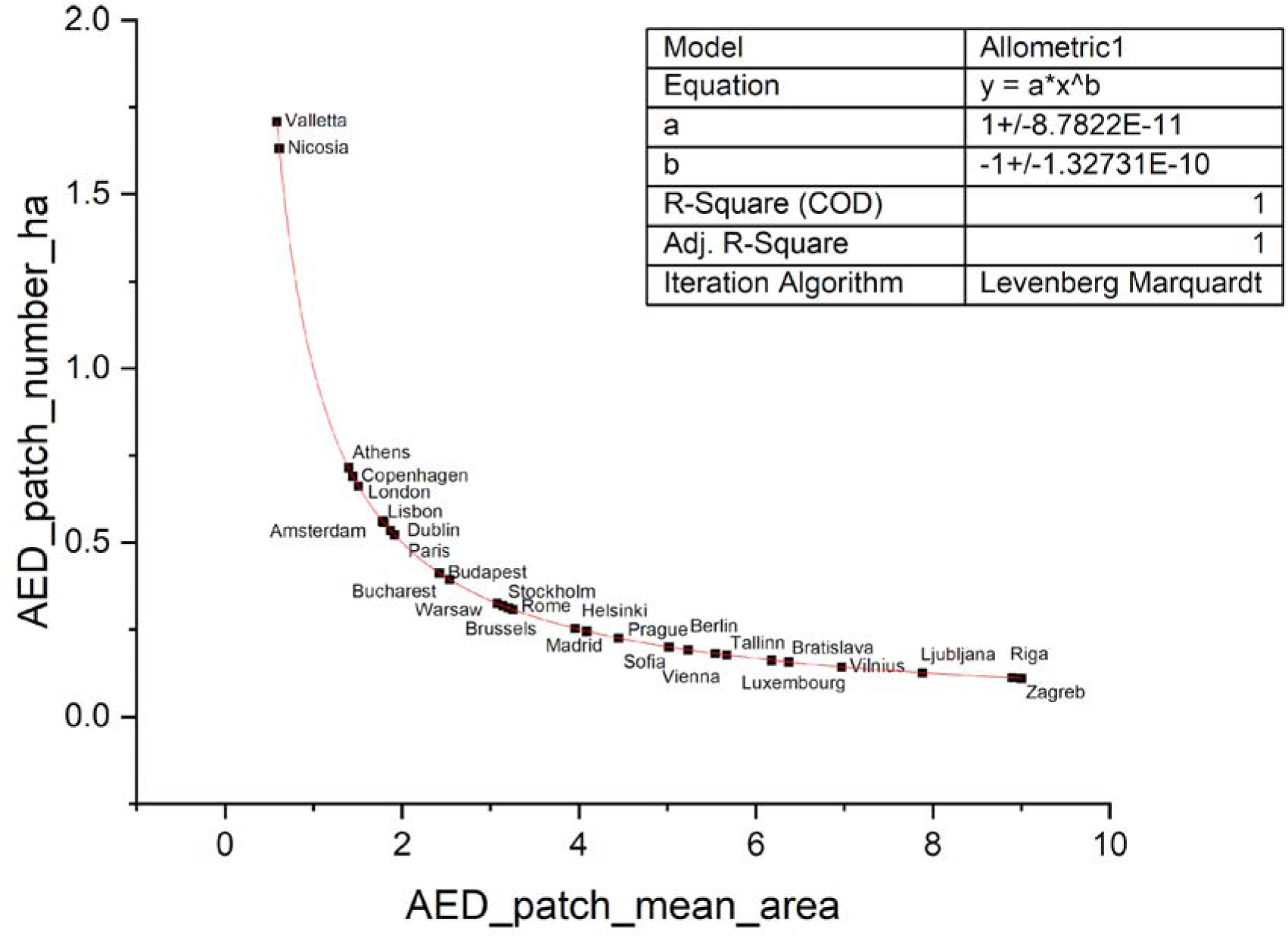
Relationship between patch mean area (*AED_patch_mean_area*) and patch number per hectare (*AED_patch_number_ha*)

Based on this relationship, we can easily detect that Nicosia and Valletta account for the lowest average dimension of the patches and the highest density in terms of the number of patches per ha. These cities also account for the lowest percentage of UGS (respectively 3% and 12% of the landscape). Many of the AED metrics are less interesting because linearly directly related to the amount of UGS, such as: *AED_patch_mean_area*, *AED_pland_trees*, *AED_gyrate*, *AED_lpi* (R^2^ of 0.63, 0.99, 0.51, 0.46 respectively), or inversely related, like for *AED_patch_number_ha* (R^2^ = 0.74).

It is interesting to note that the relationship between UGS total area (*AED_ta*), and its percentage expression against the total landscape areas is quite small (R^2^=0.18) because the strong variation in the dimension of the different Capitals makes the comparison between them very difficult.

The only metric not correlated with UGS percentage coverage is the edge density (*AED*_*ed*), which is instead directly inversely correlated to the aggregation of the patches (inverse linear decreasing relationship with *ADD_pladj*, with R^2^=0.99) and sensitive to the shape of the patches too (direct linear increasing relationship with *SHP_para_cv*, with R^2^=0.32).

Finally, the metric *AED_pland_shrubs* has limited interest because shrublands represent a very small part of the total UGS area; the average is less than 1% of the investigated areas, with notable exceptions only for Madrid (7%) and Valletta (4%), typical of a more Mediterranean condition. On average, the correlation between these metrics and the simple indicator of the percentage of area covered by UGS is good (R^2^ = 0.42).

#### 3.1.3 Core area metrics

Regarding the core area metrics the empirical analysis of the data confirmed the theoretical direct relationship between the amount of habitat available (*AED_ta*) and the amount of core areas *COR_tca* (R^2^=0.98); it is interesting to note, because less theoretical evident, that also the variability in the size of the patches (*AED_area_cv*) is directly correlated to the variability in the size of the core areas (*COR_core_cv*) with R^2^=0.99. The relationship between the percentage of the core areas on the total UGS (on the y axis) and the percentage of UGS on the total investigated areas (on the x axis, Figure 7) can easily represent the situation with Capitals poor in percentage coverage of the UGS that are also poor in core areas (R^2^ = 0.66) such as Nicosia and Valletta. The percentage of core areas is directly correlated with the average dimension of the patches (R^2^ = 0.73).

**Fig. 7.**
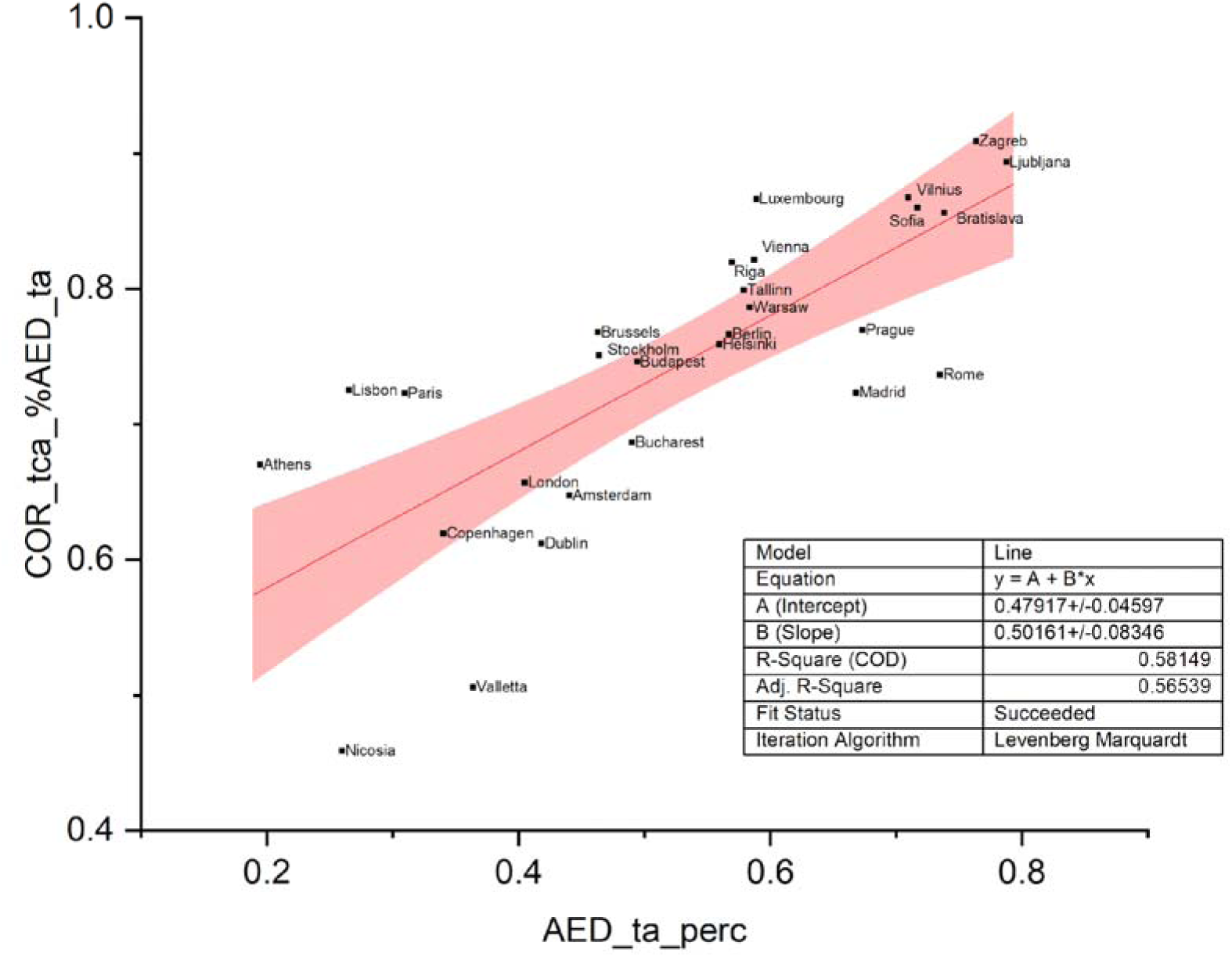
Relationship between the percentage of the core areas on the total UGS (y axis) and the percentage of UGS on the total investigated areas (x axis)

Here, the number of disjunct core areas (*COR_dcad*) is inversely related to the presence in the landscape of very large patches (*AED_lpi*, R^2^ = 0.41) but once again directly related to the percentage of UGS (R^2^=0.61). On average, the correlation between these metrics and the reference indicator percentage of area covered by UGS is weak (R^2^=0.27).

#### 3.1.4 Diversity metrics

The diversity metrics exhibited moderate collinearity (R^2^ = 0.42). This relationship is likely constrained by the thematic detail available in the EUVM, which only classified UGS into three broad classes: trees, shrubs, and herbaceous coverage. For a more meaningful diversity analysis, at least the main tree species or categories should be mapped. Nevertheless, the average correlation between these metrics and the reference indicator percentage of area covered by UGS is very poor (R^2^=0.29), while both of them are very much related with an inverse exponential rule to the UGS area expressed as a total (R^2^=0.95 for *DIV_prd* and 0.46 with *DIV_sidi*). The relationship between the total area covered by UGS and *DIV_prd* is available in the Annex, Figure 11.

#### 3.1.5 Shape metrics

The shape metrics demonstrate, as expected, good independence from the amount of UGS in the landscape, regardless of whether analyzed in absolute or relative terms. Only the variability in the patch shapes (*SHP_frac_cv, SHP_para_cv*, *SHP_shape_cv*) demonstrates a (weak) direct relationship with the percentage amount of UGS in the landscape (R^2^ of 0.22, 0.24, and 0.34, respectively). Indeed, the three types of metrics we used for accounting the patch shape complexity are linearly correlated among each other, but while the fractal dimension (*SHP_frac_mn*) has good agreement both with the perimeter/area ratio (*SHP_para_mn*) and with the shape index (*SHP_shape_mn*) (R^2^ of 0.71 and 0.79 respectively) the perimeter/area ratio has only limited agreement with the shape index (R^2^ = 0.3).

The variability of the three metrics demonstrates good agreement within them and direct weak agreement with the average dimension of the patches (R^2^ of 0.18 between *AED_ta* and *SHP_shape_cv*) and with the dimension of the largest patch (R^2^ of 0.14 between *AED_lpi* and *SHP_para_cv*). The average correlation between these metrics and the reference indicator percentage of area covered by UGS is very weak (R^2^=0.17).

#### 3.1.6 Network metrics

An example of the ecological network reconstructed with graph theory analysis using Graphab is available in Figure 8 for the city of Paris. Information for all the EU Capital cities is available in the Annex, Figure 12. Regarding the three main NET metrics calculated with Graphab, it is important to note that they are very redundant (with positive linear relationships with R^2^ ranging between a minimum of 0.76 and a maximum of 0.97). When the analysis is repeated in the rank order of the cities, the results are almost identical, regardless of the metrics used. The second information is that the relationship between these three metrics and the reference metrics *AED_pland* ranges between 0.45 and 0.57 (and between 0.62 and 0.66, when repeated based on the rank).

**Fig. 8.**
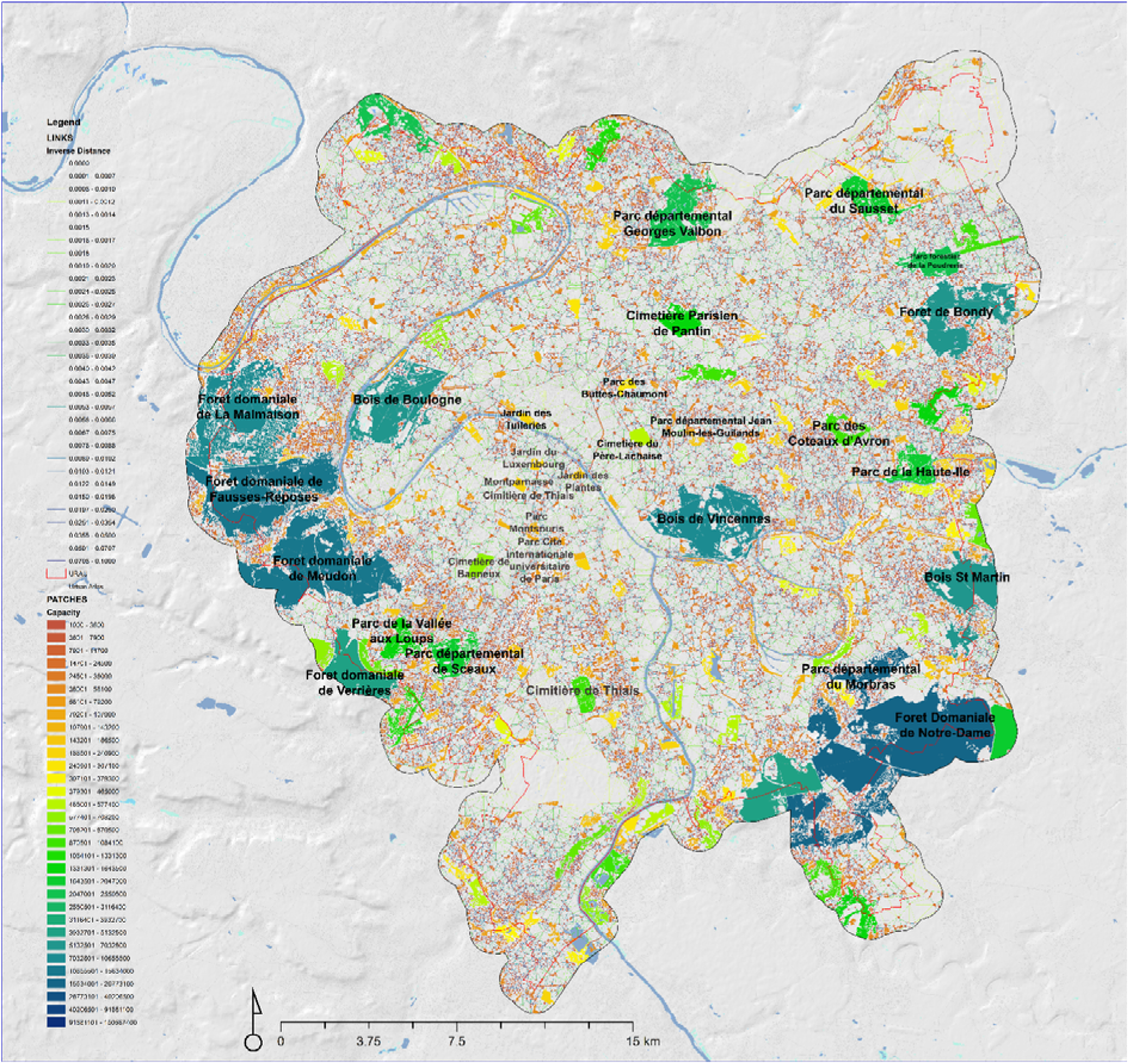
The result of the Graphab analysis, with patches and the links between them for the city of Paris. For a better understanding, some of the most important nodes of the network are labelled with their local names

Some interesting relationships were also found comparing graph-theory-based metrics and the other landscape metrics calculated with FRAGSTATS. The three NET metrics are directly correlated with the mean dimension of patches (*AED_patch_mean_area*) with R^2^ around 0.6 and inversely with *AGG_division,* also approximately with R^2^=0.6 and a direct one with *AED_lpi* and with *COR_core_cv* (R^2^ of 0.59 and 0.34, respectively).

The amount of links of the networks created in Graphab (*NET_links_mean_lenght*) is directly and strictly related to the number of patches in the area (*AED_patch_number_ha*) with R^2^ of 0.87, and inversely with the relative coverage of the vegetated area (*AED_pland*) with R^2^ of 0.68; but their average length is not related to the average size of the patches but has just a limited direct relationship with the number of patches in the area (R^2^ of 0.24) and an inverse relationship with the relative coverage of the vegetated area (R^2^ of 0.44). The variability in the length of the links (*NET_links_sd_lenght*) is not related to the variability in the dimension of the patches but is very much directly related only to the average length of the links (R^2^ of 0.68).

The Gini indicator when calculated on the patch capacity is quite redundant with the more traditional indexes related to patch density and size (inverse relationship with the density of the patches expressed by *AED_patch_number_ha* with R^2^ of 0.93 and direct relationship with the size of the patches expressed by *AED_patch_mean_area* with R^2^ of 0.68 and with relative coverage of the vegetated area expressed by *AED_pland* with R^2^ of 0.8.

While when the Gini indicator is calculated on the patch distance expressed by the links length it is very weakly related the size and shape of the patches, for example R^2^ of 0.25 with the relative coverage of the vegetated area expressed by *AED_pland* or R^2^ of 0.49 with the variability in the fractal dimension of the patches expressed by *SHP_frac_cv* but only very weakly with their average dimension (R^2^ of 0.18 with *AED_patch_mean_area*).

Finally, the last two metrics based on patch capacity (*NET_tot_patch_capacity* and *NET_Patch_mean_capacity*) are redundant with the total habitat area (*AED_ta*) and the average size of the patches (*AED_patch_mean_area*), respectively. This may be because, in this study, the capacity of each node in the network generated with Graphab is determined by the size of the patch. Overall, the correlation among the different metrics is available in the Annex (Figure 13) and the average correlation between these metrics and the reference indicator percentage of UGS is in the average (R^2^=0.45).

### 3.2 Cluster analysis and ranking

The result of the cluster analysis (Figure 9), together with those from the PCA (Figure 10), can improve the comprehension of the similarities between the analyzed cities and the role of the metrics we used for the description of UGS spatial arrangement.

**Fig. 9.**
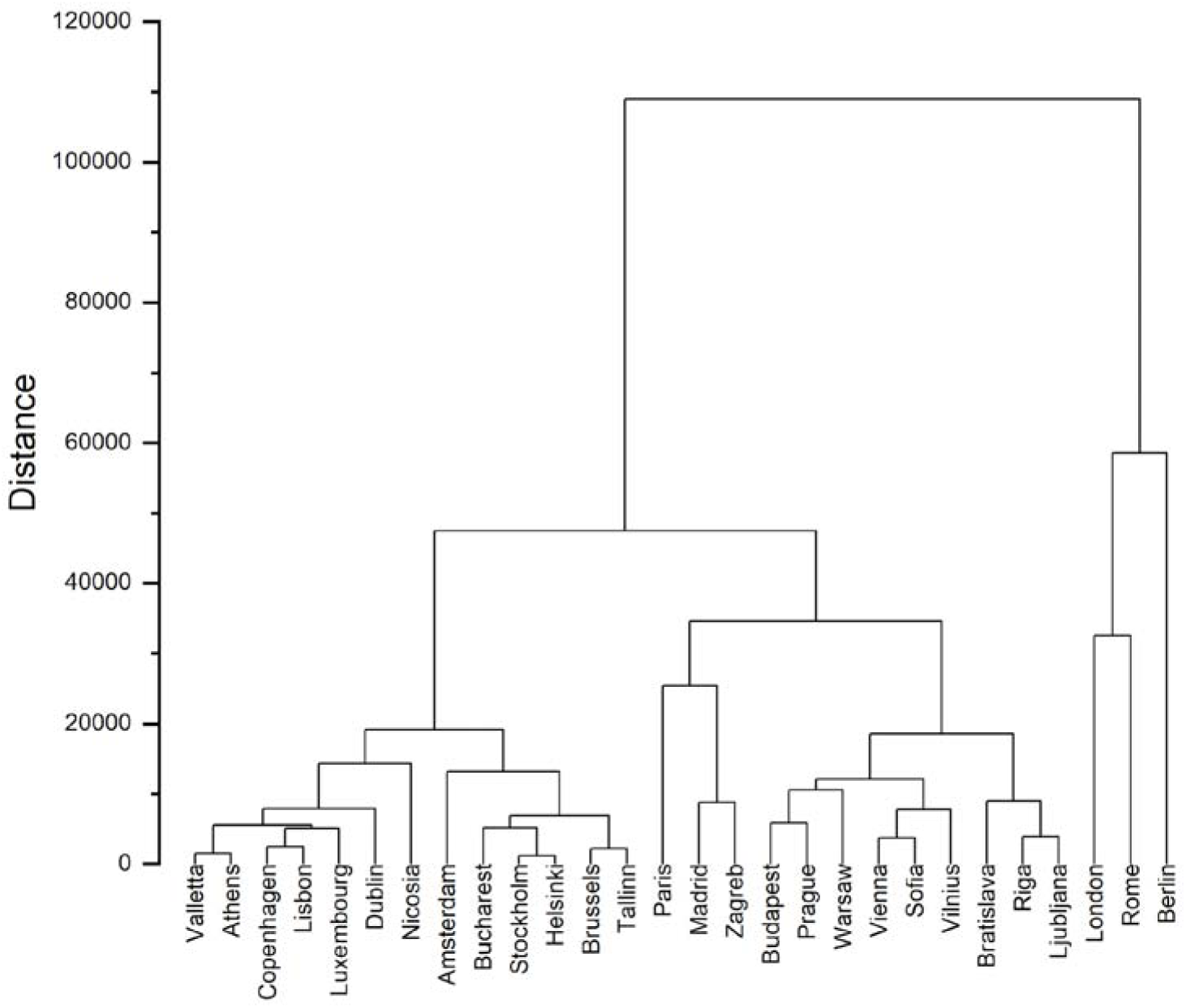
The dendrogram created with hierarchical cluster analysis on the basis of calculated metrics

**Fig. 10.**
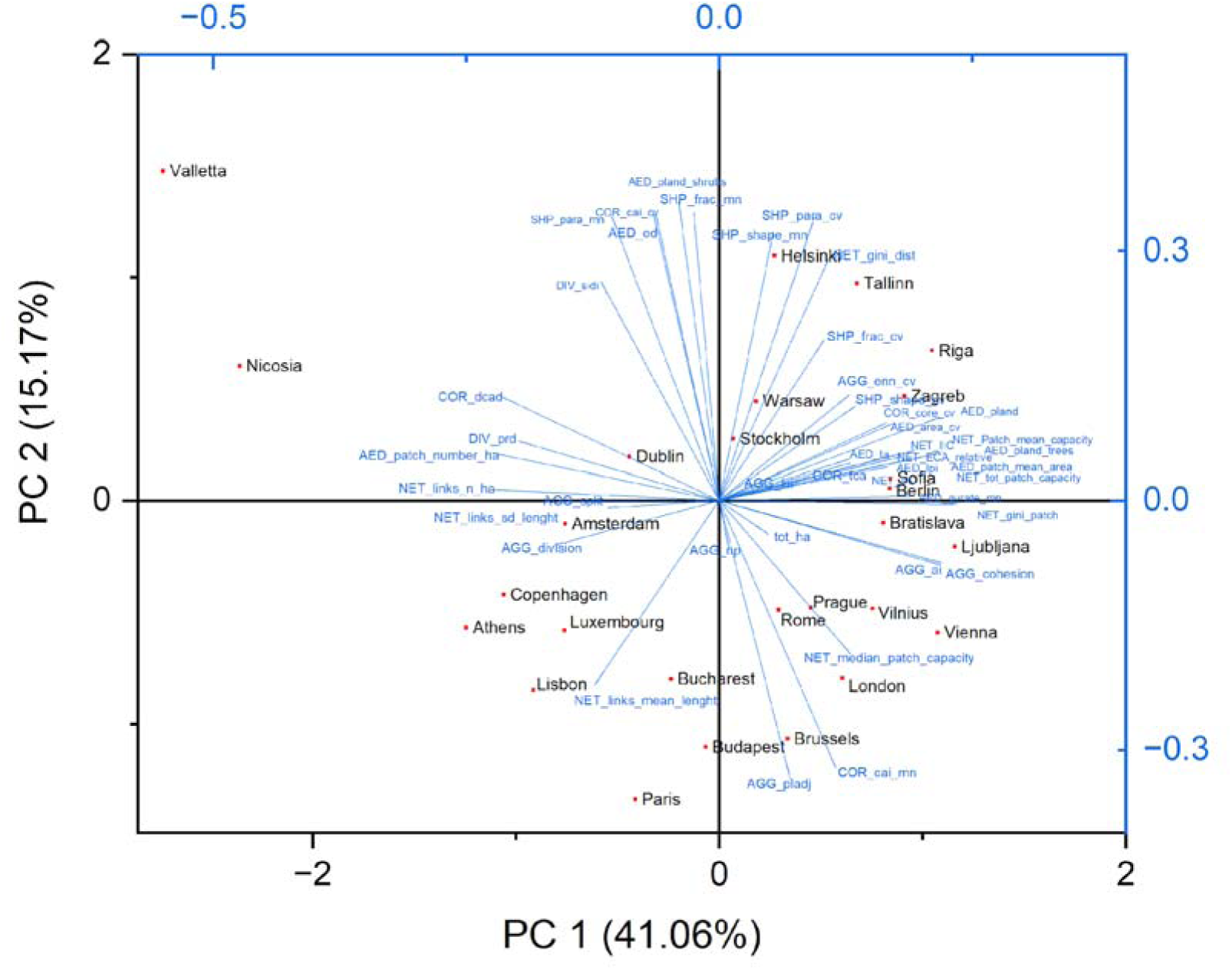
Representation of the PCA results with eigenvectors and eigenvalues for the different metrics and the position of the 28 cities in the bidimensional space created with the first two components

In Figure 9, there is an interesting ranking of the cities moving from left to right that can be interpreted as the effect of both the dimension of the areas and the spatial arrangement of UGS. Valletta and Athens are, in fact, relatively small cities, with limited UGS and weak connections in their ecological network. Another group of relatively small cities, with a broader availability of UGS than the previous group, is made up of Copenhagen, Lisbon, Nicosia, Dublin, and Luxembourg. This is also confirmed by the PCA, with all these cities resulting in the lower values of the first principal component (Figure 10).

On the right side of the dendrogram (Figure 9), we can find the largest cities (i.e., Berlin, Rome, and London). The city of Berlin has large urban forest areas distributed around the city center (the PCA shows that this condition appears to be similar to that of the city of Sofia). Moving to the left in the dendrogram we can find those cities that emerged as those with a more relevant connectivity of UGS (Ljubljana, Riga and Bratislava and then in a second cluster with Vilnius, Sofia and Vienna and a third one with Warsaw, Prague and Budapest), these cities have a smaller dimension but with a very consistent and highly connected system of relatively large forest areas in the peri-urban areas as well as green areas in the inner urban area.

The remaining cities exhibit more intermediate characteristics, both in terms of size and the connectivity of UGS, with fewer extensive forested areas adjacent to urban zones. Based on the results of the cluster analysis, specifically the sum of distances within clusters, Bratislava emerges as the most representative city, while London is the least representative.

## 4. Discussion

### 4.1. The impact of city boundary definition on comparative analysis

The first information to be considered for discussing the results we achieved is the role of the very different sizes of the cities we analyzed. They range between very large landscapes such as London and Rome (respectively 1575 and 1286 km^2^) or Berlin and Paris (respectively 892 and 800 km^2^) to very small cities such as Luxembourg, Valletta, or Athens (52, 50, and 39 km^2^ respectively). Such variability is in part due to the real differences in the built-up areas, and in part to the different approaches adopted in defining the boundaries of the urban areas in the URAU dataset. The impact of such differences on the assessment of UGS is well known (Raciti et al. 2012). Indeed, at the political level, the attempt to develop a standardized methodology for defining the boundaries of the cities is documented already (Dijkstra et al. 2019), recently leading to a new proposal based on population gridded spatial information (Eurostat 2021). Waiting for a new official revised version of these European urban areas’ boundaries, the results we achieved can be used to assess the relationship between the different metrics, but not, in our opinion, to create a conclusive rank analysis of the cities according to the different spatial arrangements of UGS.

Based on the EUVM, the mean urban vegetation percentage resulted slightly over 50% of the 28 European Capital cities we analyzed, almost half of them (13) are located below this threshold, with Athens, Nicosia, and Lisbon not reaching 30%. On the other hand, the largest share of UGS was recorded in Southeast European Capital cities – i.e., Ljubljana and Zagreb – with more than 75% of the URAU area covered by vegetation. These results are confirmed by literature. For example, in 2016, Ljubljana was awarded the “Green Capital Award” (European Commission, 2016), a European recognition for cities above 100,000 inhabitants that excel in environmental sustainability, building on their commitment to initiatives like the European Green Deal, the Zero Pollution Action Plan, the Circular Economy Action Plan, and the Biodiversity Strategy. In the past several years, Ljubljana was thoroughly studied for its features (i.e., a three-hill ridge that runs across the city (Pirnat 1997), creating a natural forest corridor) and large, managed urban forests within the city core boundaries (Jovanović and Glišić 2021).

### 4.2. Interrelationships among landscape metrics and urban classification

Given the necessary precautions regarding the impact of city dimension on the metrics, our analysis focused on identifying and interpreting the relationships among the various landscape metrics.

In detail, one of the founding motivations for this study is the assessment of the relationship between the reference indicators most frequently used for assessing and ranking green infrastructures of cities, the percentage of area covered by UGS (expressed in this study by the indicator *AED_pland*), and the other metrics related to the spatial arrangement of UGS.

Our empirical results confirmed the relationships between landscape metrics that we already know from their theoretical definition: a specific habitat that in a given landscape is very frequent is expected to be more connected, less fragmented, and with more core areas than a landscape where the same habitat is very rare (Forman 1995). In fact, in our dataset, when the area covered by UGS represents a more relevant percentage of the landscape (*AED_pland* is higher) than the density of patches (*AED_patch_number_ha*) is lower, and their average dimension (*AED_patch_mean_area*) is higher (a few big patches), and thus also the relative amount of core areas is higher (*COR_cai_mn*). But in general, the agreement between *AED_pland* and all the other tested indicators was quite weak (R^2^=0.34). The weakness is particularly true for the indicators related to the patch shape (R^2^=0.17), core areas (R^2^=0.27), diversity (R^2^=0.29), and aggregation (R^2^=0.30). The indicators related to area and edges, and those based on graph theory, denote higher correlations (R^2^ of 0.42 and 0.45, respectively).

This examination was particularly relevant given the inclusion of several metrics based on graph theory (the main ones are those expressed by *NET_PC*, *NET_ECA_relative* and *NET_IIS*).

First, our results also demonstrate that the three graph theory metrics appear extremely redundant, especially when the analysis is carried out based on the rank order; the relationship between *NET_PC* and *NET_IIS* is almost perfect. When we rank the cities based on these two metrics, the rank order for 12 cities is the same, and for the other 13, the change is only 1 position. When the ordering rank is instead compared between ECA and the other two metrics, the order changes consistently (Tallinn has a change of 8 positions, Stockholm 5, Bratislava 4, 5 cities have a change of 3 positions, 5 have 2, 7 have 1, and 7 have no changes).

In order to better understand the meaning of these three graph theory metrics, it is important to understand their relationship with the amount of UGS (*AED_pland*) in the different cities. For example, Tallinn has a relatively small part of its landscape covered by UGS, but the patches are well connected, the same, even if with a lower magnitude in Stockholm. This means that when the cities are ranked on the basis of graph-theory metrics, they may result in different ranking positions despite recording similar values in traditional landscape metrics, and vice versa. This is the case of Tallinn, which has almost double the percentage of area covered by UGS compared to Rome, ranked 3^rd^ and 20^th^ for this indicator, respectively. Still, the two cities have very similar rank positions (11^th^ and 12^th^) based on *NET_PC* and *NET_IIC*, respectively.

When we analyze the metrics related to the network links, it is important to remember that those can be created only between habitat patches. Thus, the more the area is covered by a given habitat, the less space is available for the creation of the links. Also, more links and longer links mean less connectivity inside the habitat patches. Furthermore, the global landscape connectivity metrics, computed for the urban vegetation network, highlighted the remarkable vegetation connectivity within Ljubljana, which achieved the highest values of *NET_PC* and the third highest *NET_IIC* scores among the European capital cities included in the analysis. However, as recommended by Saura and Pascual-Hortal (Saura and Pascual-Hortal 2007), the intrinsic characteristics of these metrics should be carefully considered. For instance, *NET_IIC* assigns higher values to patches at short and intermediate distances compared to *NET_PC*, impacting the overall connectivity assessment (Bodin and Saura 2010). Moreover, the potential discrepancies between probabilistic (i.e., *NET_PC*) and binary (i.e., *NET_IIC*) connectivity metrics arise from their inherent characteristics. Probabilistic metrics incorporate randomness through probability distributions, which act as weightings for decision likelihood. In contrast, binary metrics provide a deterministic view by merely indicating whether patches are connected or not. Despite the strong positive correlation between *NET_PC* and *NET_IIC* obtained in the current study, current research suggests that when confronted with conflicting results, urban landscape management should give priority to *NET_PC* over *NET_IIC* analysis, as it appears to be a more suitable approach and prevents oversimplification of patch connections (Babí Almenar et al. 2019).

Overall, based on the cluster analysis results (Figure 9), we found that three main groups of cities can be considered. A first group of relatively small cities with poor availability of UGS and limited connectivity where the limited size also limit the possibility to have in the UGS large forest peri-urban areas, followed by a group of medium-size cities with strong differences in terms of spatial arrangement of UGS but with a consistent component of peri-urban forest area and finally a small group of very large cities where the size of the urban areas determine very different conditions with a complex landscape mosaic and a vast interconnected system of UGS.

### 4.3 Opportunities and limitations

The study presents several key opportunities for urban ecological planning by demonstrating that graph-theory-based metrics, when combined with traditional landscape ecology metrics, offer a more comprehensive assessment of UGS spatial arrangement and connectivity than relying solely on the oversimplified percentage area covered by UGS. This multi-metric approach challenges the utility of the percentage of the total landscape belonging to a class (i.e., tree or shrubs) as a standalone policy indicator, given its poor correlation with most spatial metrics, and provides a standardized, scalable methodology using Earth observation data for cross-city comparison. The cluster analysis further provides an opportunity for developing context-specific planning strategies by grouping cities based on size and UGS arrangement.

However, the present analysis is constrained by the high heterogeneity of urban boundary definitions, which prevents the establishment of conclusive city rank analysis. Furthermore, the low thematic resolution of the EUVM, which only classified UGS into three broad classes, limits the efficacy of diversity metrics and suggests that future mapping must include the main tree species or additional categories for a more meaningful ecological assessment. Indeed, challenges remain in systematically mapping small-scale green spaces (e.g., single tree species, especially in private gardens), non-tree species, and understory vegetation in urban environments (Amir Reza Shahtah massebi 2018; Neyns and Canters 2022).

## 5. Conclusions

Based on this research, we can draw four main conclusions. First, remote sensing, particularly Copernicus products, provides a robust basis for standardized mapping of urban vegetation in Europe. Using Copernicus layers, we produced a 10 m European Urban Vegetation Map (EUVM) with an overall accuracy of 0.84, adequate for analyzing UGS spatial configuration in major European cities. We recommend for the future extending EUVM to all EU cities and updating it as new Copernicus products become available. The recent release of Copernicus High Resolution Layers (e.g., Cropland) will improve discrimination between herbaceous vegetation and agricultural land by incorporating landCuse information. Future studies should consider mapping of dominant urban tree species in order to obtain a more refined biodiversity assessment.

Second, the commonly used indicator “percentage of UGS” is overly simplistic for ranking or monitoring urban greening. It should be reported alongside metrics that capture patch shape, core-area extent, spatial arrangement, and compositional diversity. Although these limitations are well known in landscape ecology, this study provides the first quantitative, acrossCcapital evidence that reliance on percentCcover alone can be misleading; we therefore encourage reconsideration of its primacy in official guidance.

Third, graphCbased connectivity analysis is a valuable complement to traditional landscape metrics, revealing aspects of UGS spatial organization not captured by conventional tools and identifying candidate areas for targeted planting or planning to enhance connectivity. Graphab and the graph4lg R package allow computation of local metrics per habitat patch and spatial interpolation of results. However, graph metrics should be interpreted together with standard landscape indicators: some cities with clearly different UGS configurations can exhibit similar network metrics values, underscoring the need for a combined approach.

Finally, using the full indicator range, we produced the first standardized classification of UGS spatial configurations for all European capitals. The resulting dataset can support stakeholders and policymakers by improving visualization and comparative analysis, informing interventions to promote sustainable, biodiverse, and socially inclusive urban green infrastructure.

## Acknowledgements

This article was carried out in the framework of the URBANA project, ESA Contract No. 4000143311/23/I-DT for “DEVELOPMENT AND VERIFICATION OF URBAN ANALYTICS”, the LIFE ESCAPOS (Proposal number: 101157553) “Environment energy for Strategic CApillary urban POlicieS” and the “Advanced models for estimating carbon uptake by urban vegetation to support ecological transition policies” project, funded by the Tuscany Region under the initiative “Green Transition – Advanced Training Projects with Research Grants” (CUP B53C23002970009), within the European Social Fund Plus (ESF+) 2021–2027 Regional Programme – Priority 4 ‘Youth Employment. The following projects also support the activities: FORWARDS. H2020 project funded by the European Commission, number 101084481 call HORIZON-CL6-2022-CLIMATE-01-05. MONIFUN. H2020 project funded by the European Commission, number 101134991 call HORIZON-CL6-2023-CircBio-01-14. NextGenCarbon. H2020 project funded by the European Commission, number 101184989 call HORIZON-CL5-2024-D1-01-07. SUPERB. H2020 project funded by the European Commission, number 101036849 call LC-GD-7-1-2020; MULTIFOR “Multi-scale observations to predict Forest response to pollution and climate change” PRIN 2020 Research Project of National. Relevance funded by the Italian Ministry of University and Research (prot. 2020E52THS).

## Statements & Declarations

### 7.3 Author Contributions

Conceptualization, C.B., S.F., S.M., L.C., G.C, J.M, G.D.L., G.C.; Data curation, C.B., S.F., G.C.; Formal analysis, C.B.; Investigation, C.B., L.T., G.C., G.C., Methodology, C.B., G.C.; Project administration, S.M., G.C., G.C.; Writing—original draft, C.B. Writing—review and editing, C.B., S.F., L.C., S.M., L.T., G.C., J.M, E.V, G.D.A, G.D.L., G.C. All authors have read and agreed to the published version of the manuscript.

### 7.4 Data Availability

The data that support the findings of this study are available from the corresponding author upon reasonable request.

## Annex

**Fig. 11.**
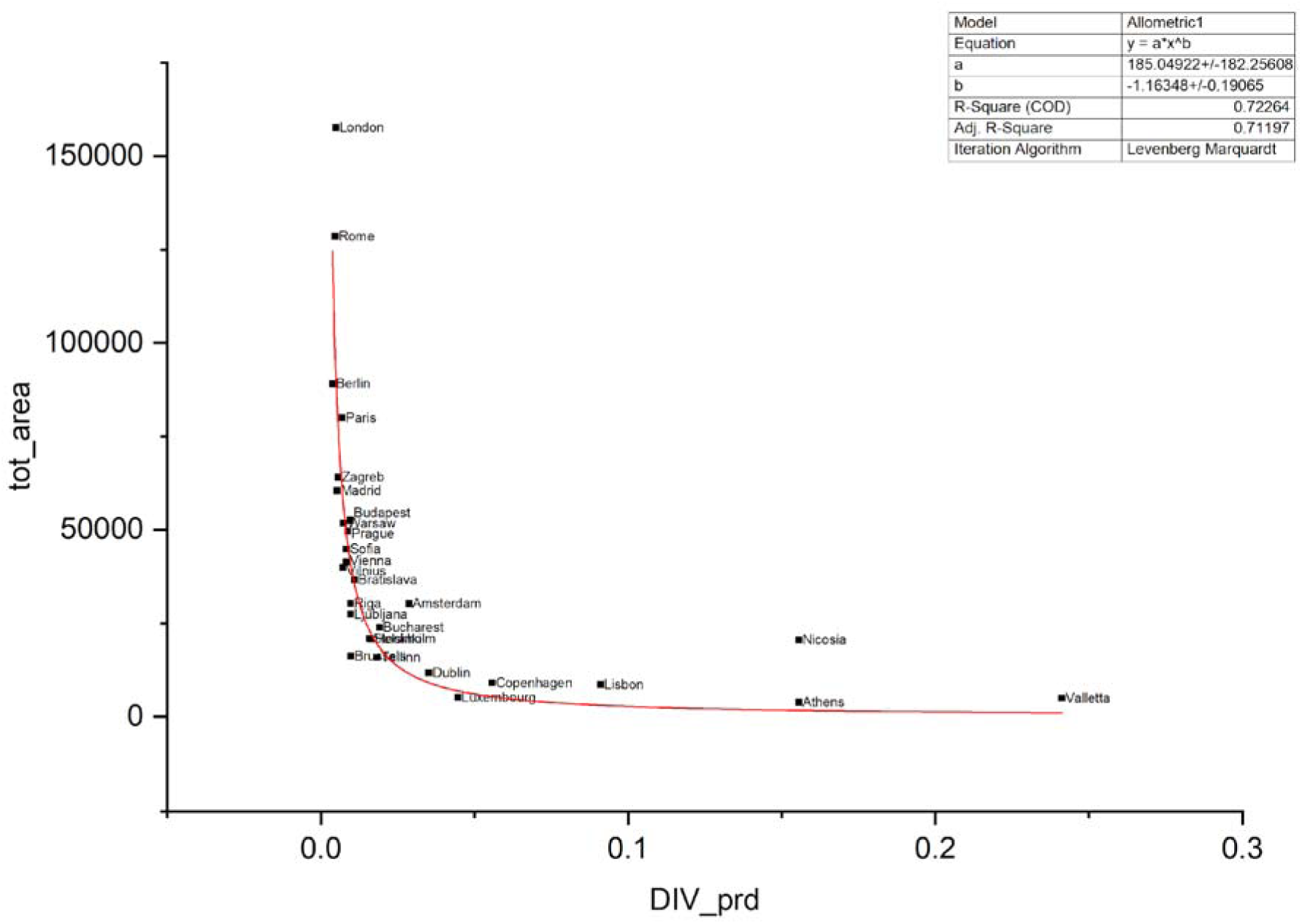
The relationship between the total area covered by the area of UGS (tot_area) and DIV_prd.

**Fig. 12.**
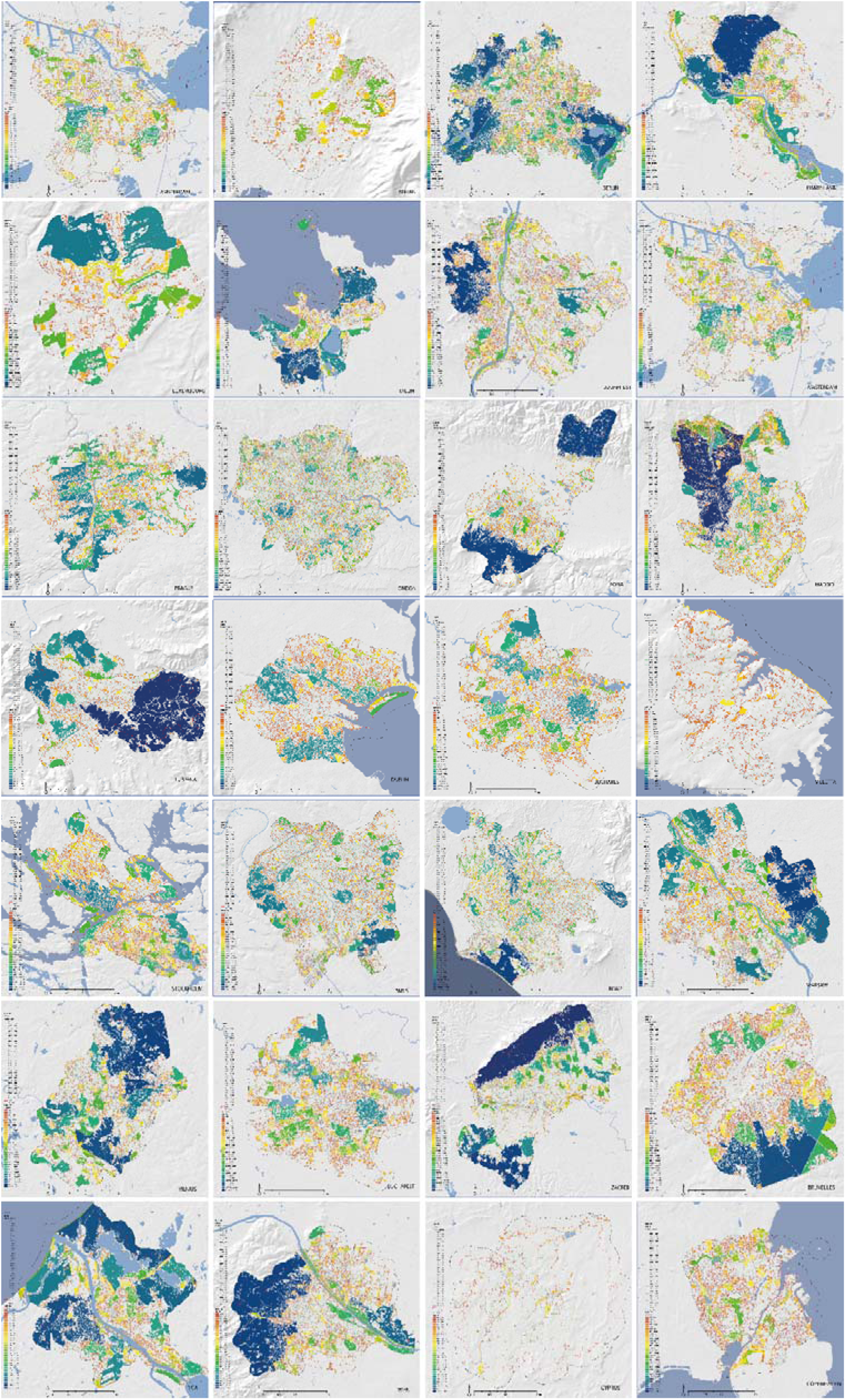
The result of the Graphab analysis for the 28 investigated European capitals with patches and the links between them

**Fig. 13.**
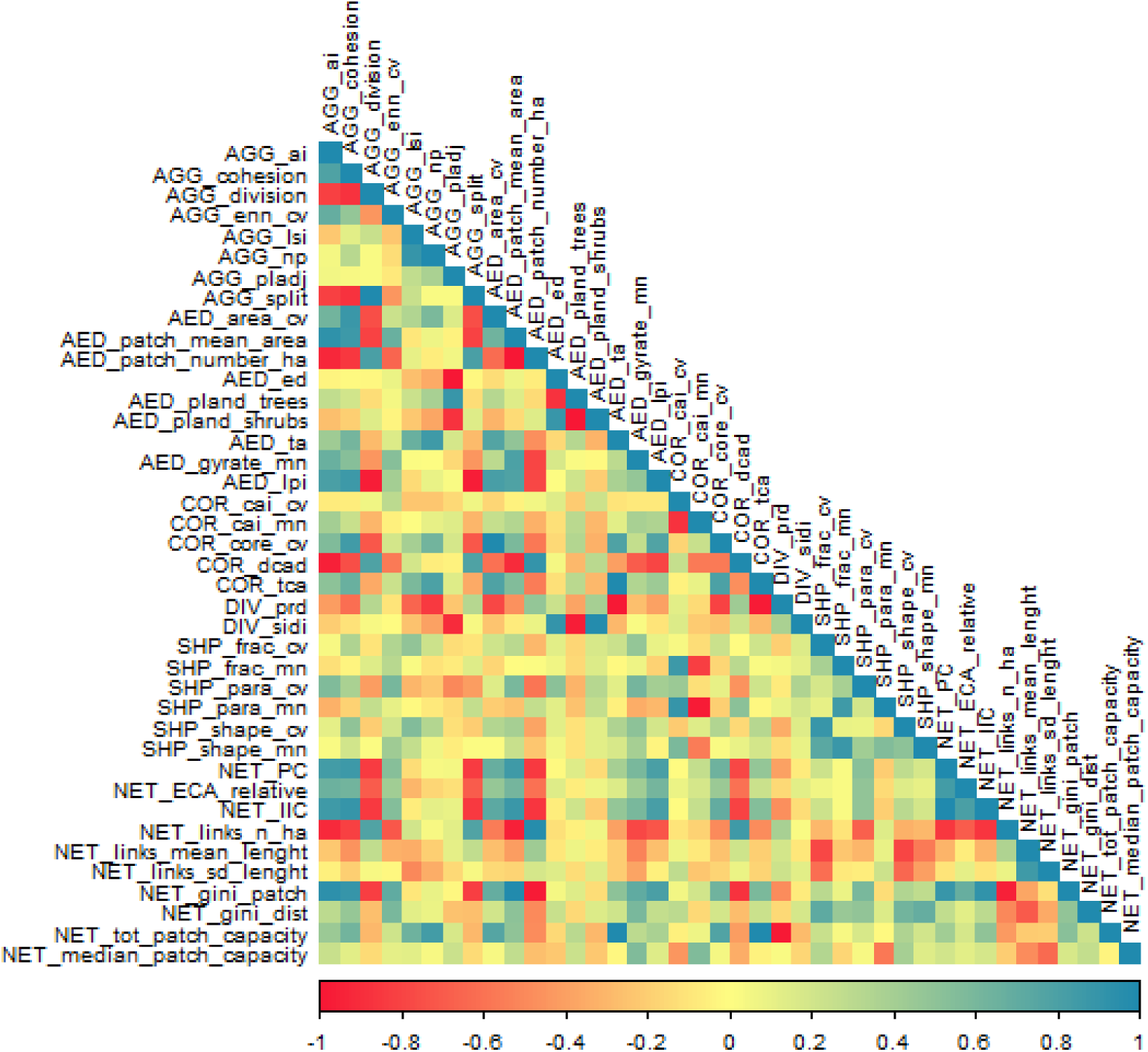
Correlation among different landscape metrics (as reported in Table 2A)

---

**Table 1A.** Validation results for each Country’s capital city, considering a 1km boundary. Overall Accuracy (OA), Producer’s accuracy (PA), and User’s Accurcay (UA) are reported as long as the LUCAS sampled points falling within each boundary (LUCAS_pt) and the number of True Positive (TP), True Negative (TN), False Positive (FP) and False Negative (FN).

| City | OA | PA | UA | LUCAS_pt | TP (n°) | TN (n°) | FP (n°) | FN (n°) |
| --- | --- | --- | --- | --- | --- | --- | --- | --- |
| Amsterdam | 0.89 | 0.40 | 0.80 | 110 | 31 | 62 | 4 | 13 |
| Athens | 1.00 | 0.00 | 0.00 | 29 | 6 | 15 | 0 | 8 |
| Berlin | 0.85 | 0.67 | 0.93 | 58 | 13 | 39 | 5 | 1 |
| Bratislava | 0.89 | 0.81 | 0.89 | 83 | 1 | 69 | 13 | 0 |
| Bruxelles | 0.72 | 0.43 | 1.00 | 73 | 15 | 44 | 2 | 12 |
| Bucharest | 0.81 | 0.37 | 0.88 | 106 | 28 | 62 | 2 | 14 |
| Budapest | 0.79 | 0.50 | 0.70 | 11 | 1 | 8 | 0 | 2 |
| Copenhagen | 0.82 | 0.33 | 1.00 | 21 | 4 | 12 | 1 | 4 |
| Dublin | 0.64 | 0.11 | 1.00 | 11 | 0 | 11 | 0 | 0 |
| Helsinki | 0.87 | 0.85 | 0.91 | 144 | 37 | 68 | 10 | 29 |
| Lisbon | 1.00 | 1.00 | 1.00 | 60 | 29 | 23 | 3 | 5 |
| Ljubljana | 0.92 | 0.83 | 0.94 | 119 | 27 | 68 | 8 | 16 |
| London | 0.85 | 0.46 | 0.70 | 86 | 34 | 43 | 6 | 3 |
| Luxembourg | 0.77 | 0.50 | 0.67 | 48 | 7 | 31 | 3 | 7 |
| Madrid | 0.73 | 0.56 | 0.79 | 22 | 1 | 13 | 0 | 8 |
| Nicosia | 0.84 | 1.00 | 0.07 | 164 | 25 | 113 | 7 | 19 |
| Paris | 0.80 | 0.63 | 0.77 | 76 | 27 | 41 | 4 | 4 |
| Prague | 0.81 | 0.56 | 0.88 | 13 | 2 | 8 | 1 | 2 |
| Riga | 0.79 | 0.56 | 0.93 | 58 | 14 | 32 | 1 | 11 |
| Rome | 0.84 | 0.57 | 0.78 | 28 | 3 | 22 | 0 | 3 |
| Sofia | 0.90 | 0.93 | 0.72 | 61 | 4 | 50 | 1 | 6 |
| Stockholm | 0.88 | 0.77 | 0.92 | 53 | 18 | 27 | 0 | 8 |
| Tallinn | 0.76 | 0.50 | 0.80 | 14 | 2 | 12 | 0 | 0 |
| Valletta | 0.89 | 0.50 | 1.00 | 67 | 7 | 47 | 1 | 12 |
| Vilnius | 0.89 | 0.87 | 0.87 | 73 | 24 | 40 | 2 | 7 |
| Warsaw | 0.85 | 0.69 | 1.00 | 49 | 15 | 30 | 1 | 3 |
| Wien | 0.85 | 0.70 | 0.89 | 53 | 17 | 30 | 2 | 4 |
| Zagreb | 0.90 | 0.92 | 0.85 | 176 | 16 | 134 | 7 | 19 |

**Table 2A.**
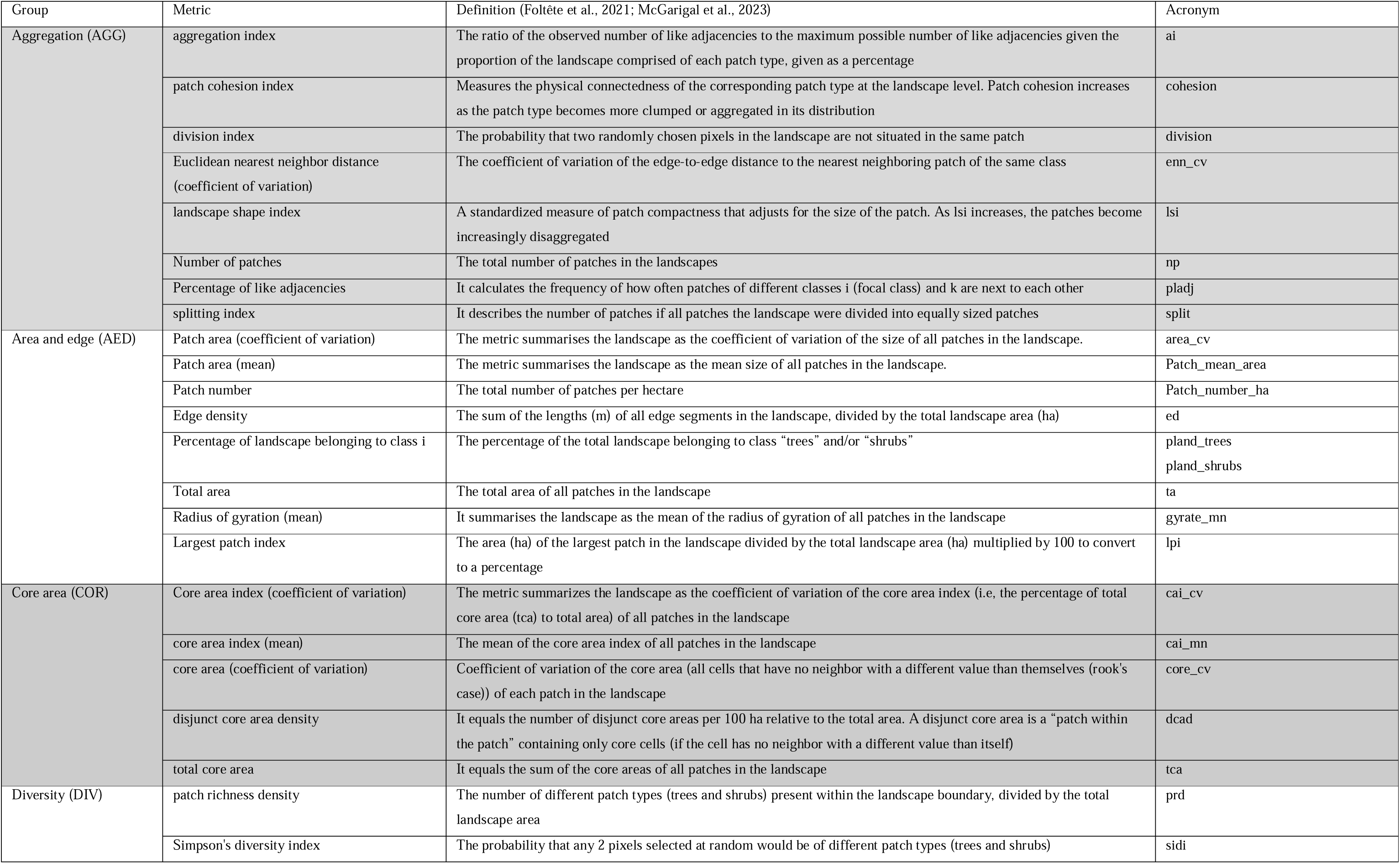
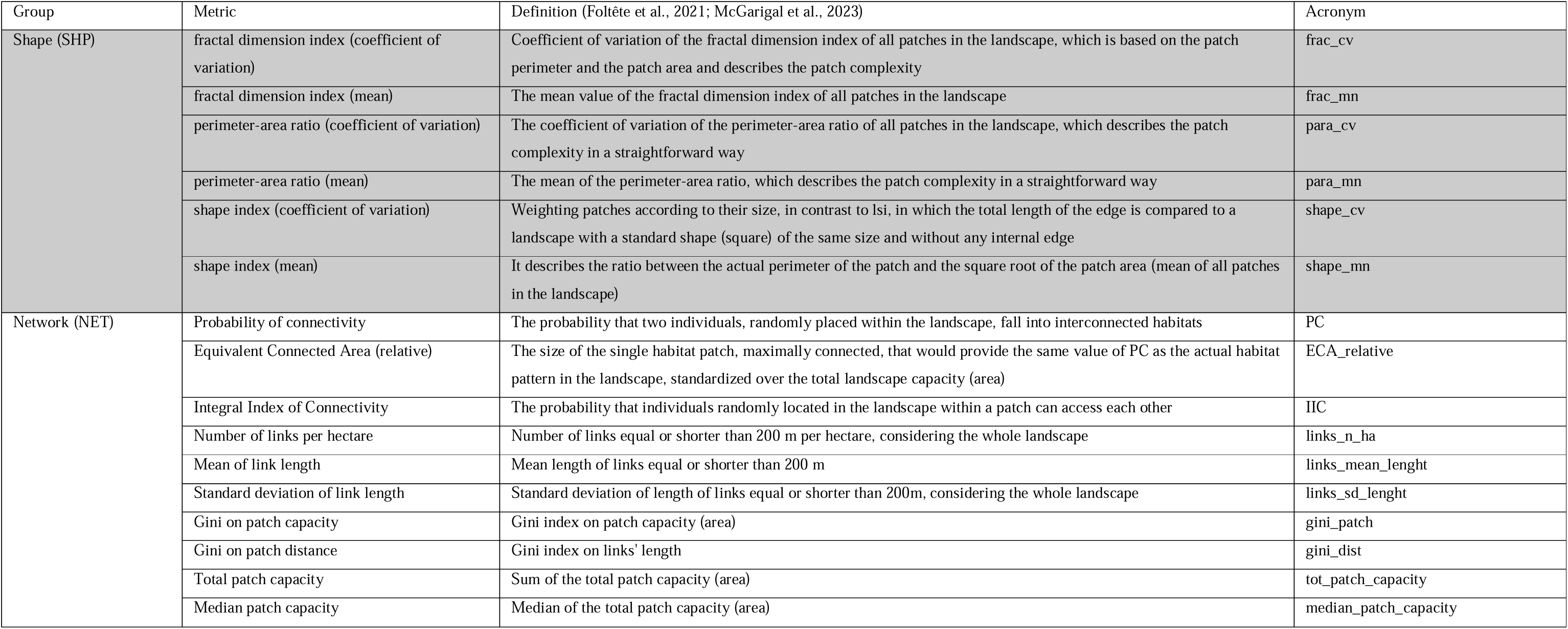
Landscape patterns and connectivity metrics calculated for each city.

**Table 3A.** Urban land cover based on the European Map of Vegetation, calculated for each city as an absolute value (in km^2^) and percentage of the total area. The categories are non-vegetated areas (Non Veg), trees, shrubs, grass, and total vegetation (Veg) as the sum of trees, shrubs and grass.

| City | Non Veg (km<sup>2</sup>) | Trees (km<sup>2</sup>) | Shrubs (km<sup>2</sup>) | Grass (km<sup>2</sup>) | Tot. area (km<sup>2</sup>) | Veg (km<sup>2</sup>) | Non Veg (%) | Trees (%) | Shrubs (%) | Grass (%) | Veg (%) |
| --- | --- | --- | --- | --- | --- | --- | --- | --- | --- | --- | --- |
| Amsterdam | 169.77 | 60.59 | 0.38 | 72.67 | 303.4 | 133.64 | 55.95 | 19.97 | 0.13 | 23.95 | 44.05 |
| Athens | 31.41 | 7.34 | 0 | 0.24 | 39 | 7.59 | 80.54 | 18.83 | 0 | 0.63 | 19.46 |
| Berlin | 386.04 | 432.74 | 1.59 | 71.49 | 891.86 | 505.82 | 43.28 | 48.52 | 0.18 | 8.02 | 56.72 |
| Bratislava | 96.2 | 146.2 | 1.53 | 123.58 | 367.51 | 271.31 | 26.18 | 39.78 | 0.42 | 33.63 | 73.83 |
| Brussels | 87.25 | 69.57 | 0 | 5.65 | 162.47 | 75.22 | 53.7 | 42.82 | 0 | 3.48 | 46.3 |
| Bucharest | 122.17 | 79.35 | 0.67 | 37.38 | 239.57 | 117.4 | 51 | 33.12 | 0.28 | 15.6 | 49.01 |
| Budapest | 265.71 | 165.16 | 0.21 | 94.11 | 525.19 | 259.48 | 50.59 | 31.45 | 0.04 | 17.92 | 49.41 |
| Copenhagen | 59.91 | 20.17 | 1.72 | 8.96 | 90.76 | 30.85 | 66.01 | 22.23 | 1.89 | 9.87 | 33.99 |
| Dublin | 68.49 | 39.24 | 2.4 | 7.56 | 117.68 | 49.19 | 58.2 | 33.34 | 2.04 | 6.42 | 41.8 |
| Helsinki | 91.81 | 95.37 | 0.41 | 20.83 | 208.42 | 116.61 | 44.05 | 45.76 | 0.2 | 9.99 | 55.95 |
| Lisbon | 63.53 | 17.91 | 0.58 | 4.46 | 86.47 | 22.94 | 73.47 | 20.71 | 0.67 | 5.16 | 26.53 |
| Ljubljana | 58.44 | 142.49 | 0.9 | 73.22 | 275.06 | 216.61 | 21.25 | 51.81 | 0.33 | 26.62 | 78.75 |
| London | 937.73 | 364.97 | 0.66 | 272.37 | 1575.73 | 637.99 | 59.51 | 23.16 | 0.04 | 17.29 | 40.49 |
| Luxembourg | 21.28 | 20.85 | 0.23 | 9.38 | 51.74 | 30.46 | 41.13 | 40.3 | 0.44 | 18.13 | 58.87 |
| Madrid | 200.78 | 262.63 | 42.16 | 99.43 | 604.99 | 404.21 | 33.19 | 43.41 | 6.97 | 16.43 | 66.81 |
| Nicosia | 152.63 | 6.5 | 0 | 47.02 | 206.14 | 53.51 | 74.04 | 3.15 | 0 | 22.81 | 25.96 |
| Paris | 552.91 | 210.85 | 0.01 | 36.57 | 800.34 | 247.43 | 69.08 | 26.35 | 0 | 4.57 | 30.92 |
| Prague | 162.22 | 179.7 | 1.42 | 153.12 | 496.46 | 334.24 | 32.68 | 36.2 | 0.29 | 30.84 | 67.32 |
| Riga | 130.96 | 146.87 | 2.51 | 23.82 | 304.15 | 173.19 | 43.06 | 48.29 | 0.82 | 7.83 | 56.94 |
| Rome | 340.95 | 346.7 | 3.14 | 595.17 | 1285.96 | 945.01 | 26.51 | 26.96 | 0.24 | 46.28 | 73.49 |
| Sofia | 127.39 | 179.18 | 0.06 | 143.39 | 450.02 | 322.63 | 28.31 | 39.82 | 0.01 | 31.86 | 71.69 |
| Stockholm | 112.58 | 85.45 | 0.07 | 11.89 | 209.98 | 97.4 | 53.61 | 40.7 | 0.03 | 5.66 | 46.39 |
| Tallinn | 67.2 | 79.06 | 1.1 | 12.11 | 159.46 | 92.27 | 42.14 | 49.58 | 0.69 | 7.59 | 57.86 |
| Valletta | 31.97 | 4 | 2.03 | 12.23 | 50.23 | 18.26 | 63.64 | 7.96 | 4.05 | 24.35 | 36.36 |
| Vienna | 171.28 | 179.03 | 0.12 | 64.41 | 414.84 | 243.56 | 41.29 | 43.16 | 0.03 | 15.53 | 58.71 |
| Vilnius | 116.39 | 197.55 | 0.66 | 85.98 | 400.58 | 284.19 | 29.06 | 49.32 | 0.17 | 21.46 | 70.95 |
| Warsaw | 215.48 | 218.81 | 0.41 | 82.53 | 517.23 | 301.75 | 41.66 | 42.3 | 0.08 | 15.96 | 58.34 |
| Zagreb | 151.66 | 273.15 | 2.39 | 214.09 | 641.3 | 489.63 | 23.65 | 42.59 | 0.37 | 33.38 | 76.35 |

**Table 4A.** Aggregation metrics (AGG) calculated in the 28 European Capitals.

| City | AGG_ai | AGG_cohesion | AGG_division | AGG_enn_cv | AGG_lsi | AGG_np | AGG_pladj | AGG_split |
| --- | --- | --- | --- | --- | --- | --- | --- | --- |
| Amsterdam | 87.35 | 98.26 | 0.98 | 117.7 | 106.45 | 3893 | 99.94 | 62.19 |
| Athens | 88.02 | 97.32 | 0.97 | 112.32 | 43.72 | 919 | 99.82 | 37.73 |
| Berlin | 91.98 | 99.69 | 0.94 | 250.66 | 186.51 | 10310 | 99.94 | 17.13 |
| Bratislava | 95.02 | 99.49 | 0.83 | 305.81 | 67.43 | 2868 | 99.85 | 6.06 |
| Brussels | 91.95 | 99.53 | 0.83 | 75.95 | 82.48 | 3159 | 100 | 6.05 |
| Bucharest | 89.28 | 99.1 | 0.97 | 214.43 | 110.37 | 4130 | 99.92 | 31.02 |
| Budapest | 90.97 | 99.34 | 0.92 | 90.19 | 131.79 | 8680 | 99.98 | 12.52 |
| Copenhagen | 86.58 | 97.15 | 0.99 | 105.87 | 79.98 | 2482 | 99.45 | 109.58 |
| Dublin | 86.07 | 99.19 | 0.94 | 81.73 | 106 | 3045 | 99.97 | 15.69 |
| Helsinki | 91.2 | 98.85 | 0.96 | 437.48 | 98.15 | 3101 | 99.94 | 28.29 |
| Lisbon | 90.53 | 98.02 | 0.92 | 131.91 | 44.92 | 1233 | 99.73 | 13.05 |
| Ljubljana | 96.34 | 99.8 | 0.67 | 265.91 | 53.26 | 2641 | 99.91 | 3 |
| London | 87.46 | 98.24 | 1 | 105.24 | 255.24 | 27276 | 99.99 | 233.68 |
| Luxembourg | 95.37 | 99.24 | 0.87 | 270.63 | 31.8 | 726 | 99.94 | 7.93 |
| Madrid | 90.38 | 99.8 | 0.8 | 151.4 | 172.45 | 9347 | 98.13 | 5.03 |
| Nicosia | 79.14 | 93.76 | 0.99 | 180.7 | 53.57 | 1048 | 100 | 83.81 |
| Paris | 90.09 | 98.88 | 0.98 | 85.69 | 169.42 | 15093 | 100 | 40.08 |
| Prague | 92.09 | 99.59 | 0.93 | 228.13 | 120.32 | 5167 | 99.91 | 14.39 |
| Riga | 93.76 | 99.73 | 0.9 | 266.91 | 89.9 | 2336 | 99.85 | 10.38 |
| Rome | 90.81 | 99.42 | 0.95 | 140.53 | 191.23 | 13519 | 99.9 | 19.61 |
| Sofia | 95.04 | 99.74 | 0.74 | 322.42 | 78.13 | 4833 | 99.99 | 3.8 |
| Stockholm | 90.95 | 98.89 | 0.98 | 177.09 | 102.73 | 4040 | 99.98 | 44.81 |
| Tallinn | 92.93 | 99.73 | 0.82 | 414.71 | 74.25 | 1926 | 99.87 | 5.43 |
| Valletta | 81.48 | 92.38 | 0.99 | 118.95 | 52.49 | 1417 | 98.33 | 140.15 |
| Vienna | 93.9 | 99.83 | 0.68 | 246.85 | 96.14 | 4394 | 99.99 | 3.11 |
| Vilnius | 95.37 | 99.7 | 0.88 | 252.02 | 78.34 | 4031 | 99.97 | 8.57 |
| Warsaw | 92.23 | 99.34 | 0.94 | 273.08 | 129.41 | 8903 | 99.98 | 16.31 |
| Zagreb | 96.79 | 99.73 | 0.79 | 207.62 | 60.28 | 3959 | 99.86 | 4.7 |

**Table 5A.** Area and Edge metrics (AED) calculated in the 28 European Capitals.

| City | AED_area_cv | AED_patch_mean_area | AED_patch_number_ha | AED_ed | AED_pland_trees | AED_pland_shrubs | AED_ta | AED_gyrate_mn | AED_lpi |
| --- | --- | --- | --- | --- | --- | --- | --- | --- | --- |
| Amsterdam | 784.98 | 1.8 | 0.56 | 1.03 | 0.2 | 0 | 6991.58 | 43.1 | 9.1 |
| Athens | 483.56 | 1.4 | 0.72 | 3.11 | 0.19 | 0 | 1285.1 | 36.29 | 7.55 |
| Berlin | 2451.61 | 5.23 | 0.19 | 1.11 | 0.49 | 0 | 53967.02 | 47.34 | 17.54 |
| Bratislava | 2173.58 | 6.37 | 0.16 | 2.81 | 0.4 | 0 | 18269.68 | 54.17 | 38.15 |
| Brussels | 2283.22 | 3.25 | 0.31 | 0 | 0.43 | 0 | 10276.97 | 44.52 | 39.05 |
| Bucharest | 1149.62 | 2.54 | 0.39 | 1.47 | 0.33 | 0 | 10491.44 | 42.85 | 10.86 |
| Budapest | 2631.35 | 2.42 | 0.41 | 0.37 | 0.31 | 0 | 21040.66 | 42.32 | 27.28 |
| Copenhagen | 465.4 | 1.45 | 0.69 | 9.45 | 0.22 | 0.02 | 3590.16 | 42.3 | 3.41 |
| Dublin | 1389.9 | 1.87 | 0.53 | 0.54 | 0.33 | 0.02 | 5695.3 | 41.55 | 15.94 |
| Helsinki | 1042.44 | 3.96 | 0.25 | 1.02 | 0.46 | 0 | 12274.73 | 52.76 | 15.78 |
| Lisbon | 967.27 | 1.78 | 0.56 | 4.89 | 0.21 | 0.01 | 2197.46 | 40.53 | 22.36 |
| Ljubljana | 2964.91 | 7.88 | 0.13 | 1.7 | 0.52 | 0 | 20823.81 | 46.03 | 56.13 |
| London | 1075.77 | 1.51 | 0.66 | 0.18 | 0.23 | 0 | 41169.81 | 39.31 | 3.46 |
| Luxembourg | 952.1 | 6.18 | 0.16 | 1.03 | 0.4 | 0 | 4484.74 | 55.52 | 28.59 |
| Madrid | 4310.62 | 4.09 | 0.24 | 34.33 | 0.43 | 0.07 | 38227.11 | 45.32 | 44.28 |
| Nicosia | 339.35 | 0.61 | 1.63 | 0 | 0.03 | 0 | 642.69 | 34.47 | 6.06 |
| Paris | 1938 | 1.92 | 0.52 | 0.01 | 0.26 | 0 | 28930.91 | 39.17 | 11.26 |
| Prague | 1892.41 | 4.45 | 0.22 | 1.69 | 0.36 | 0 | 22996.02 | 48.84 | 22.49 |
| Riga | 1497.22 | 8.9 | 0.11 | 2.84 | 0.48 | 0.01 | 20782.7 | 54.92 | 19 |
| Rome | 2623.64 | 3.2 | 0.31 | 1.73 | 0.27 | 0 | 43303.55 | 48.91 | 20.12 |
| Sofia | 3566.51 | 5.02 | 0.2 | 0.15 | 0.4 | 0 | 24253.57 | 40.39 | 38.1 |
| Stockholm | 944.36 | 3.14 | 0.32 | 0.32 | 0.41 | 0 | 12686.39 | 49.43 | 10.41 |
| Tallinn | 1881.65 | 5.67 | 0.18 | 2.44 | 0.5 | 0.01 | 10919.45 | 46.63 | 34.93 |
| Valletta | 301.94 | 0.59 | 1.71 | 27.56 | 0.08 | 0.04 | 829.32 | 30.87 | 4.2 |
| Vienna | 3757.39 | 5.54 | 0.18 | 0.16 | 0.43 | 0 | 24350.02 | 47.51 | 55.69 |
| Vilnius | 2167.04 | 6.97 | 0.14 | 0.62 | 0.49 | 0 | 28094.66 | 47.38 | 28.48 |
| Warsaw | 2334.43 | 3.07 | 0.33 | 0.31 | 0.42 | 0 | 27368.06 | 40.77 | 21.51 |
| Zagreb | 2899.57 | 9 | 0.11 | 2.77 | 0.43 | 0 | 35643.88 | 49.85 | 40.53 |

**Table 6A.** Core area (COR) and diversity (DIV) metrics calculated in the 28 European Capitals.

| City | COR_cai_cv | COR_cai_mn | COR_core_cv | COR_dcad | COR_tca | DIV_prd | DIV_sidi |
| --- | --- | --- | --- | --- | --- | --- | --- |
| Amsterdam | 46.85 | 33.94 | 933.77 | 150.8 | 4528.28 | 0.03 | 0.01 |
| Athens | 47.03 | 35.86 | 559.99 | 158.2 | 861.38 | 0.16 | 0.08 |
| Berlin | 36.97 | 40.3 | 2768.54 | 86.23 | 41359.85 | 0 | 0.01 |
| Bratislava | 37.39 | 43.16 | 2417.79 | 37.02 | 15640.9 | 0.01 | 0.02 |
| Brussels | 32.77 | 44.55 | 2587.35 | 66.88 | 7890.67 | 0.01 | 0 |
| Bucharest | 32.31 | 42.75 | 1309.38 | 95.94 | 7205.14 | 0.02 | 0.02 |
| Budapest | 31.49 | 43.35 | 3077.78 | 80.54 | 15707.24 | 0.01 | 0 |
| Copenhagen | 32.23 | 41.23 | 554.7 | 137.99 | 2223.93 | 0.06 | 0.09 |
| Dublin | 41.02 | 34.19 | 1596.34 | 182.94 | 3485.62 | 0.04 | 0.09 |
| Helsinki | 59.98 | 33.98 | 1244.79 | 96.2 | 9319.92 | 0.02 | 0.01 |
| Lisbon | 40.57 | 40.79 | 1180.48 | 100.62 | 1594.11 | 0.09 | 0.13 |
| Ljubljana | 40.49 | 39.55 | 3136.36 | 28.01 | 18611.43 | 0.01 | 0.01 |
| London | 46.26 | 34.49 | 1329.66 | 145.45 | 27054.44 | 0 | 0 |
| Luxembourg | 36.19 | 45.7 | 1027.1 | 31.51 | 3885.77 | 0.04 | 0.01 |
| Madrid | 59.56 | 31.1 | 4910.04 | 115.87 | 27659.92 | 0.01 | 0.27 |
| Nicosia | 66.81 | 24.67 | 483.73 | 348.54 | 294.9 | 0.16 | 0 |
| Paris | 30.62 | 42.35 | 2369.86 | 101.37 | 20918.63 | 0.01 | 0 |
| Prague | 33.77 | 43.2 | 2070.96 | 51.83 | 17693.87 | 0.01 | 0.01 |
| Riga | 41.33 | 39.04 | 1584.08 | 54.85 | 17031.45 | 0.01 | 0.03 |
| Rome | 38.03 | 40.81 | 3188.05 | 73.98 | 31909.45 | 0 | 0.02 |
| Sofia | 47.4 | 35.6 | 3886.42 | 44.19 | 20855.4 | 0.01 | 0 |
| Stockholm | 51.82 | 36.95 | 1043.75 | 89.1 | 9529.31 | 0.02 | 0 |
| Tallinn | 52.62 | 34.36 | 1998.25 | 72.2 | 8724.25 | 0.02 | 0.02 |
| Valletta | 57.76 | 29.58 | 442.66 | 281.07 | 419.65 | 0.24 | 0.45 |
| Vienna | 32.58 | 43.76 | 4092.91 | 43.66 | 20003.58 | 0.01 | 0 |
| Vilnius | 38.84 | 41.29 | 2309.5 | 35.43 | 24369.54 | 0.01 | 0.01 |
| Warsaw | 57.37 | 31.14 | 2692.34 | 90.18 | 21515.5 | 0.01 | 0 |
| Zagreb | 62.11 | 34.91 | 3073.01 | 23.49 | 32407.99 | 0.01 | 0.02 |

**Table 7A.** Shape (SHP) metrics calculated in the 28 European Capitals.

| City | SHP_frac_cv | SHP_frac_mn | SHP_para_cv | SHP_para_mn | SHP_shape_cv | SHP_shape_mn |
| --- | --- | --- | --- | --- | --- | --- |
| London | 4.55829 | 1.099304 | 29.37656 | 0.105548 | 48.22349 | 1.620655 |
| Rome | 5.150586 | 1.092584 | 30.01283 | 0.089112 | 64.67979 | 1.648261 |
| Berlin | 5.351137 | 1.095962 | 29.48761 | 0.090317 | 83.5647 | 1.700307 |
| Paris | 4.519565 | 1.083671 | 25.21146 | 0.088327 | 46.22801 | 1.526005 |
| Zagreb | 4.311407 | 1.105608 | 38.83555 | 0.104971 | 40.01334 | 1.659287 |
| Madrid | 5.682219 | 1.11429 | 52.53453 | 0.117068 | 79.31366 | 1.820272 |
| Budapest | 4.714273 | 1.082724 | 27.61176 | 0.086604 | 50.18052 | 1.538545 |
| Warsaw | 4.834903 | 1.113004 | 31.40116 | 0.112245 | 47.56531 | 1.726225 |
| Prague | 5.073451 | 1.085507 | 31.15479 | 0.085513 | 72.07524 | 1.608124 |
| Sofia | 4.720795 | 1.096329 | 30.69161 | 0.102381 | 47.80409 | 1.610705 |
| Vienna | 4.938421 | 1.085153 | 28.17366 | 0.084293 | 63.59218 | 1.585153 |
| Vilnius | 4.648738 | 1.08743 | 31.15439 | 0.089988 | 53.3317 | 1.572157 |
| Bratislava | 5.235227 | 1.093287 | 33.8654 | 0.086953 | 56.27718 | 1.660243 |
| Riga | 5.374511 | 1.106101 | 38.01724 | 0.095199 | 84.94071 | 1.792029 |
| Amsterdam | 5.281038 | 1.103448 | 28.61367 | 0.10464 | 56.36764 | 1.69678 |
| Ljubljana | 4.780971 | 1.087387 | 33.21776 | 0.09205 | 48.89887 | 1.561911 |
| Bucharest | 5.154363 | 1.081109 | 27.82255 | 0.08627 | 70.08946 | 1.572379 |
| Stockholm | 5.061944 | 1.109843 | 35.52912 | 0.101912 | 54.06514 | 1.758397 |
| Helsinki | 5.422711 | 1.128828 | 37.08175 | 0.109385 | 57.94745 | 1.943617 |
| Nicosia | 4.641932 | 1.120589 | 27.31582 | 0.12654 | 35.36626 | 1.709601 |
| Brussels | 4.620311 | 1.084916 | 29.31235 | 0.083795 | 55.09956 | 1.561532 |
| Tallinn | 5.199241 | 1.112635 | 32.68229 | 0.107607 | 70.17051 | 1.786362 |
| Dublin | 5.320235 | 1.106855 | 26.54715 | 0.104948 | 69.88132 | 1.729443 |
| Copenhagen | 4.916929 | 1.09064 | 30.02082 | 0.089645 | 48.92141 | 1.598796 |
| Lisbon | 4.289724 | 1.084375 | 32.21691 | 0.089194 | 35.94261 | 1.514516 |
| Luxembourg | 4.553345 | 1.088218 | 36.74546 | 0.083304 | 43.15458 | 1.586448 |
| Valletta | 4.093757 | 1.098221 | 29.649 | 0.114914 | 28.00114 | 1.541401 |
| Athens | 4.505419 | 1.093161 | 30.2926 | 0.103701 | 44.50046 | 1.567807 |

**Table 8A.** Network (NET) metrics calculated in the 28 European Capitals.

| City | NET_PC | NET_ECA_rel<br>ative | NET_IIC | NET_links_n_<br>ha | NET_links_me<br>an_lenght | NET_links_sd_<br>lenght | NET_gini_patc<br>h | NET_gini_dist | NET_tot_patc<br>h_capacity | NET_median_<br>patch_capacity |
| --- | --- | --- | --- | --- | --- | --- | --- | --- | --- | --- |
| Amsterdam | 0 | 0.1 | 0 | 1.25 | 69.98 | 54.12 | 0.83 | 0.6 | 7008.72 | 0.28 |
| Athens | 0.01 | 0.38 | 0.01 | 1.32 | 79.88 | 58.27 | 0.8 | 0.49 | 1290.14 | 0.26 |
| Berlin | 0.02 | 0.31 | 0.02 | 0.72 | 54.4 | 48.32 | 0.92 | 0.6 | 51153.99 | 0.28 |
| Bratislava | 0.03 | 0.37 | 0.02 | 0.47 | 59.56 | 51.44 | 0.93 | 0.67 | 17574.5 | 0.32 |
| Brussels | 0.05 | 0.49 | 0.04 | 1.04 | 64.37 | 49.56 | 0.88 | 0.53 | 9665.55 | 0.32 |
| Bucharest | 0.01 | 0.23 | 0.01 | 1.32 | 65.26 | 52.56 | 0.84 | 0.59 | 9680.22 | 0.28 |
| Budapest | 0.01 | 0.22 | 0.01 | 1.35 | 68.96 | 52.52 | 0.85 | 0.54 | 19473.74 | 0.28 |
| Copenhagen | 0 | 0.2 | 0 | 2.32 | 67.33 | 51.45 | 0.75 | 0.56 | 3208.19 | 0.29 |
| Dublin | 0.03 | 0.39 | 0.02 | 1.37 | 62.77 | 50.98 | 0.84 | 0.55 | 5704.45 | 0.29 |
| Helsinki | 0.02 | 0.39 | 0.01 | 0.65 | 60.64 | 50.99 | 0.9 | 0.63 | 12298.49 | 0.31 |
| Lisbon | 0.01 | 0.16 | 0 | 1.45 | 78.27 | 58.36 | 0.81 | 0.52 | 2037.23 | 0.26 |
| Ljubljana | 0.13 | 0.82 | 0.1 | 0.34 | 73.6 | 56.81 | 0.96 | 0.55 | 20304.92 | 0.26 |
| London | 0 | 0.08 | 0 | 1.63 | 74.62 | 53.45 | 0.81 | 0.52 | 41429.48 | 0.28 |
| Luxembourg | 0.1 | 0.87 | 0.06 | 0.45 | 70.46 | 53.73 | 0.93 | 0.55 | 4312.68 | 0.33 |
| Madrid | 0.06 | 0.47 | 0.04 | 0.75 | 46.72 | 51.56 | 0.91 | 0.69 | 36404.75 | 0.24 |
| Nicosia | 0 | 0.01 | 0 | 2.46 | 84.66 | 59.8 | 0.65 | 0.56 | 656.78 | 0.2 |
| Paris | 0.01 | 0.18 | 0.01 | 1.66 | 74.48 | 53.7 | 0.83 | 0.52 | 26625.9 | 0.27 |
| Prague | 0.02 | 0.26 | 0.01 | 0.76 | 56.89 | 49.77 | 0.9 | 0.64 | 21484.93 | 0.31 |
| Riga | 0.11 | 0.77 | 0.08 | 0.41 | 58.95 | 52.53 | 0.95 | 0.62 | 20003.12 | 0.28 |
| Rome | 0.03 | 0.43 | 0.03 | 0.92 | 74.1 | 56.68 | 0.88 | 0.57 | 40200.73 | 0.28 |
| Sofia | 0.09 | 0.9 | 0.08 | 0.51 | 69.94 | 54.29 | 0.94 | 0.62 | 24218.79 | 0.28 |
| Stockholm | 0.02 | 0.46 | 0.02 | 0.93 | 54.71 | 47.58 | 0.87 | 0.63 | 12707.58 | 0.31 |
| Tallinn | 0.03 | 0.78 | 0.03 | 0.46 | 60.47 | 52.95 | 0.94 | 0.69 | 10911.03 | 0.28 |
| Valletta | 0 | 0.07 | 0 | 3.77 | 70.9 | 58.84 | 0.62 | 0.56 | 847.92 | 0.22 |
| Vienna | 0.08 | 0.55 | 0.06 | 0.58 | 63.83 | 52.98 | 0.92 | 0.59 | 23392.74 | 0.32 |
| Vilnius | 0.06 | 0.6 | 0.05 | 0.42 | 70.55 | 54.99 | 0.95 | 0.56 | 27173.43 | 0.25 |
| Warsaw | 0.05 | 0.43 | 0.04 | 0.94 | 65.47 | 51.78 | 0.9 | 0.55 | 27378.63 | 0.26 |
| Zagreb | 0.05 | 0.58 | 0.05 | 0.2 | 73.92 | 57.58 | 0.96 | 0.59 | 35666.69 | 0.27 |

## References

Afrifa, J. K., Monney, K. A., & Deikumah, J. P. (2022). Effects of urban land-use types on avifauna assemblage in a rapidly developing urban settlement in Ghana. Urban Ecosystems. 10.1007/s11252-022-01281-0

Amir Reza Shahtah massebi. (2018). Remote sensing of urban green spaces: A review | Elsevier Enhanced Reader. 1996, 1–37. https://reader.elsevier.com/reader/sd/pii/S1618866720307639?token=7370115A9B6B7EA7E051361AE43B5A6E855B807D07945069E6AFF8D0ABC0E7D717274841554C599D5281E994A08B2AED&originRegion=eu-west-1&originCreation=20210621165700

Babí Almenar, J., Bolowich, A., Elliot, T., Geneletti, D., Sonnemann, G., & Rugani, B. (2019). Assessing habitat loss, fragmentation and ecological connectivity in Luxembourg to support spatial planning. Landscape and Urban Planning, 189(May), 335–351. 10.1016/j.landurbplan.2019.05.004

Ballin, M., Barcaroli, G., Masselli, M., & Scarnó, M. (2018). Redesign sample for Land Use/Cover Area frame Survey (LUCAS) 2018.

Bodin, Ö., & Saura, S. (2010). Ranking individual habitat patches as connectivity providers: Integrating network analysis and patch removal experiments. Ecological Modelling, 221(19), 2393–2405. 10.1016/j.ecolmodel.2010.06.017

Brandt, J., Ertel, J., Spore, J., & Stolle, F. (2023). Wall-to-wall mapping of tree extent in the tropics with Sentinel-1 and Sentinel-2. Remote Sensing of Environment, 292, 113574. 10.1016/j.rse.2023.113574

Bulman, C. R., Wilson, R. J., Holt, A. R., Bravo, L. G., Early, R. I., Warren, M. S., & Thomas, C. D. (2007). MINIMUM VIABLE METAPOPULATION SIZE, EXTINCTION DEBT, AND THE CONSERVATION OF A DECLINING SPECIES. Ecological Applications, 17(5), 1460–1473. 10.1890/06-1032.1

Chang-Fu, L., Bin, Z., Xing-Yuan, H., & Wei, C. (2010). Selection of distance thresholds of urban forest landscape connectivity in Shenyang City. Yingyong Shengtai Xuebao, 21(10).

Chen, Y., & Yu, S. (2017). Impacts of urban landscape patterns on urban thermal variations in Guangzhou, China. International Journal of Applied Earth Observation and Geoinformation, 54, 65–71. 10.1016/j.jag.2016.09.007

d’Andrimont, R., Yordanov, M., Martinez-Sanchez, L., Eiselt, B., Palmieri, A., Dominici, P., Gallego, J., Reuter, H. I., Joebges, C., Lemoine, G., & van der Velde, M. (2020). Harmonised LUCAS in-situ land cover and use database for field surveys from 2006 to 2018 in the European Union. Scientific Data, 7(1), 352. 10.1038/s41597-020-00675-z

Das, D. N., Chakraborti, S., Saha, G., Banerjee, A., & Singh, D. (2020). Analysing the dynamic relationship of land surface temperature and landuse pattern: A city level analysis of two climatic regions in India. City and Environment Interactions, 8, 100046. 10.1016/j.cacint.2020.100046

Dash, J., & Ogutu, B. O. (2016). Recent advances in space-borne optical remote sensing systems for monitoring global terrestrial ecosystems. Progress in Physical Geography: Earth and Environment, 40(2), 322–351. 10.1177/0309133316639403

Du, C., Ge, S., Song, P., Jombach, S., Fekete, A., & Valánszki, I. (2025). Optimizing Urban Green Spaces for Vegetation-Based Carbon Sequestration: The Role of Landscape Spatial Structure in Zhengzhou Parks, China. Forests, 16(4), 679. 10.3390/f16040679

Duncan, J. M. A., Boruff, B., Saunders, A., Sun, Q., Hurley, J., & Amati, M. (2019). Turning down the heat: An enhanced understanding of the relationship between urban vegetation and surface temperature at the city scale. Science of The Total Environment, 656, 118–128. 10.1016/j.scitotenv.2018.11.223

ESA. (2018a). Copernicus HRL: Tree Cover Density. https://land.copernicus.eu/pan-european/high-resolution-layers/forests/tree-cover-density/status-maps/tree-cover-density-2018

ESA. (2018b). Copernicus Urban Atlas. https://land.copernicus.eu/local/urban-atlas/urban-atlas-2018

EU Copernicus. (2018). Small Woody Features metadata. https://land.copernicus.eu/pan-european/high-resolution-layers/small-woody-features/small-woody-features-2018?tab=metadata

European Commission. (2020). EEA geospatial data catalogue Grassland 2018 (raster 10 m), Europe, 3-yearly, Aug. 2020. 2018.

FAO. (2000). Global Forest Resources Assessment 2000, FRA 2000 Main Report. Forestry Paper, 140, 479.

Foltête, J.-C., Clauzel, C., & Vuidel, G. (2012). A software tool dedicated to the modelling of landscape networks. Environmental Modelling & Software, 38, 316–327. 10.1016/j.envsoft.2012.07.002

Foltête, J.-C., Vuidel, G., Savary, P., Clauzel, C., Sahraoui, Y., Girardet, X., & Bourgeois, M. (2021). Graphab: An application for modeling and managing ecological habitat networks. Software Impacts, 8, 100065. 10.1016/j.simpa.2021.100065

Francini, S., Chirici, G., Chiesi, L., Costa, P., Caldarelli, G., & Mancuso, S. (2024). Global spatial assessment of potential for new peri-urban forests to combat climate change. Nature Cities, 1(4), 286–294. 10.1038/s44284-024-00049-1

Fraucqueur, L., Morin, N., Masse, A., Remy, P.-Y., Hugé, J., Kenner, C., Dazin, F., Desclée, B., & Sannier, C. (2019). A new Copernicus high resolution layer at pan-European scale: small woody features. In C. M. Neale & A. Maltese (Eds.), Remote Sensing for Agriculture, Ecosystems, and Hydrology XXI (p. 37). SPIE. 10.1117/12.2532853

Frazier, A. E., & Kedron, P. (2017). Landscape Metrics: Past Progress and Future Directions. Current Landscape Ecology Reports, 2(3), 63–72. 10.1007/s40823-017-0026-0

Gallego, J., & Bamps, C. (2008). Using CORINE land cover and the point survey LUCAS for area estimation. International Journal of Applied Earth Observation and Geoinformation, 10(4), 467–475. 10.1016/j.jag.2007.11.001

Gorelick, N., Hancher, M., Dixon, M., Ilyushchenko, S., Thau, D., & Moore, R. (2017). Google Earth Engine: Planetary-scale geospatial analysis for everyone. Remote Sensing of Environment, 202, 18–27. 10.1016/j.rse.2017.06.031

Gu, D., Andreev, K., & E. Dupre, M. (2021). Major Trends in Population Growth Around the World. China CDC Weekly, 3(28), 604–613. 10.46234/ccdcw2021.160

Han, L., Zhang, R., Wang, J., & Cao, S.-J. (2024). Spatial synergistic effect of urban green space ecosystem on air pollution and heat island effect. Urban Climate, 55, 101940. 10.1016/j.uclim.2024.101940

Hesselbarth, M. H. K., Sciaini, M., Nowosad, J., & Hanss, S. (2024). Package ‘landscapemetrics’ Landscape Metrics for Categorical Map Patterns.

Hesselbarth, M. H. K., Sciaini, M., With, K. A., Wiegand, K., & Nowosad, J. (2019). landscapemetricslJ: an open source R tool to calculate landscape metrics. Ecography, 42(10), 1648–1657. 10.1111/ecog.04617

Jutz, S., & Milagro-Pérez, M. P. (2020). Copernicus: the European Earth Observation programme. Revista de Teledetección, 56. 10.4995/raet.2020.14346

Konijnendijk, C. C. (2023). Evidence-based guidelines for greener, healthier, more resilient neighbourhoods: Introducing the 3–30–300 rule. Journal of Forestry Research, 34(3), 821–830. 10.1007/s11676-022-01523-z

Kupfer, J. A. (2012). Landscape ecology and biogeography: Rethinking landscape metrics in a post-FRAGSTATS landscape. Progress in Physical Geography, 36(3), 400–420. 10.1177/0309133312439594

McGarigal, K., Cushman, S. A., & Ene, E. (2023). FRAGSTATS v4: Spatial pattern analysis program for categorical maps. https://www.fragstats.org/

McGarigal, K., & Marks, B. J. (1995). FRAGSTATS: spatial pattern analysis program for quantifying landscape structure. In General Technical Report - US Department of Agriculture, Forest Service (Vol. 351, Issue PNW-GTR-351). US Department of Agriculture, Forest Service, Pacific Northwest Research Station.

Mu, B., Tian, G., Xin, G., Hu, M., Yang, P., Wang, Y., Xie, H., Mayer, A. L., & Zhang, Y. (2021). Measuring Dynamic Changes in the Spatial Pattern and Connectivity of Surface Waters Based on Landscape and Graph Metrics: A Case Study of Henan Province in Central China. Land, 10(5), 471. 10.3390/land10050471

Neyns, R., & Canters, F. (2022). Mapping of Urban Vegetation with High-Resolution Remote Sensing: A Review. Remote Sensing, 14(4), 1031. 10.3390/rs14041031

Norton, B. A., Coutts, A. M., Livesley, S. J., Harris, R. J., Hunter, A. M., & Williams, N. S. G. (2015). Planning for cooler cities: A framework to prioritise green infrastructure to mitigate high temperatures in urban landscapes. Landscape and Urban Planning, 134, 127–138. 10.1016/j.landurbplan.2014.10.018

Pascual-Hortal, L., & Saura, S. (2006). Comparison and development of new graph-based landscape connectivity indices: towards the priorization of habitat patches and corridors for conservation. Landscape Ecology, 21(7), 959–967. 10.1007/s10980-006-0013-z

Saura, S., Estreguil, C., Mouton, C., & Rodríguez-Freire, M. (2011). Network analysis to assess landscape connectivity trends: Application to European forests (1990–2000). Ecological Indicators, 11(2), 407–416. 10.1016/j.ecolind.2010.06.011

Saura, S., & Pascual-Hortal, L. (2007). A new habitat availability index to integrate connectivity in landscape conservation planning: Comparison with existing indices and application to a case study. Landscape and Urban Planning, 83(2–3), 91–103. 10.1016/j.landurbplan.2007.03.005

Savary, P., Foltête, J., Moal, H., Vuidel, G., & Garnier, S. (2021). graph4lg: A package for constructing and analysing graphs for landscape genetics in R. Methods in Ecology and Evolution, 12(3), 539–547. 10.1111/2041-210X.13530

Savary, P., Tannier, C., Foltête, J.-C., Bourgeois, M., Vuidel, G., Khimoun, A., Moal, H., & Garnier, S. (2024). How does dispersal shape the genetic patterns of animal populations in European cities? A simulation approach. Peer Community Journal, 4, e40. 10.24072/pcjournal.407

Schindler, S., von Wehrden, H., Poirazidis, K., Hochachka, W. M., Wrbka, T., & Kati, V. (2015). Performance of methods to select landscape metrics for modelling species richness. Ecological Modelling, 295, 107–112. 10.1016/j.ecolmodel.2014.05.012

Stroud, S., Peacock, J., & Hassall, C. (2022). Vegetation-based ecosystem service delivery in urban landscapes: A systematic review. Basic and Applied Ecology, 61, 82–101. 10.1016/j.baae.2022.02.007

United Nations. (2018). The World’s Cities in 2018. World Urbanization Prospects: The 2018 Revision, 34.

Uuemaa, E., Antrop, M., Roosaare, J., Marja, R., & Mander, Ü. (2009). Landscape metrics and indices: An overview of their use in landscape research. Living Reviews in Landscape Research, 3, 1–28. 10.12942/lrlr-2009-1

Uuemaa, E., Mander, Ü., & Marja, R. (2013). Trends in the use of landscape spatial metrics as landscape indicators: A review. Ecological Indicators, 28, 100–106. 10.1016/j.ecolind.2012.07.018

Villegas, P., Gili, T., & Caldarelli, G. (2021). Emergent spatial patterns of coexistence in species-rich plant communities. Physical Review E, 104(3), 034305. 10.1103/PhysRevE.104.034305

Villegas, P., Gili, T., Caldarelli, G., & Gabrielli, A. (2024). Evidence of scale-free clusters of vegetation in tropical rainforests. Physical Review E, 109(4), L042402. 10.1103/PhysRevE.109.L042402

Wulder, M. A., Loveland, T. R., Roy, D. P., Crawford, C. J., Masek, J. G., Woodcock, C. E., Allen, R. G., Anderson, M. C., Belward, A. S., Cohen, W. B., Dwyer, J., Erb, A., Gao, F., Griffiths, P., Helder, D., Hermosilla, T., Hipple, J. D., Hostert, P., Hughes, M. J., … Zhu, Z. (2019). Current status of Landsat program, science, and applications. Remote Sensing of Environment, 225(March), 127–147. 10.1016/j.rse.2019.02.015

Yan, J., Zhou, W., Han, L., & Qian, Y. (2018). Mapping vegetation functional types in urban areas with WorldView-2 imagery: Integrating object-based classification with phenology. Urban Forestry & Urban Greening, 31, 230–240. 10.1016/j.ufug.2018.01.021

